# Multi range ERK responses shape the proliferative trajectory of single cells following oncogene induced senescence

**DOI:** 10.1101/2022.10.06.511142

**Authors:** Jia-Yun Chen, Clemens Hug, José Reyes, Chengzhe Tian, Luca Gerosa, Fabian Fröhlich, Bas Ponsioen, Hugo J G Snippert, Sabrina L. Spencer, Ashwini Jambhekar, Peter K. Sorger, Galit Lahav

**Affiliations:** Laboratory of Systems Pharmacology, Harvard Medical School, Boston, MA 02115, USA; Department of Systems Biology, Harvard Medical School, Boston, MA 02115, USA; Department of Biochemistry, University of Colorado Boulder, Boulder, CO 80303, USA; BioFrontiers Institute, University of Colorado Boulder, Boulder, CO 80303, USA; Molecular Cancer Research, Center for Molecular Medicine, University Medical Center Utrecht, Utrecht University, The Netherlands. Oncode Institute, The Netherlands

## Abstract

Oncogene-induced senescence (OIS) is a phenomenon in which aberrant oncogene expression causes non-transformed cells to enter a non-proliferative state. Cells undergoing OIS display phenotypic heterogeneity, with some cells senescing and others remaining proliferative. The causes of the heterogeneity remain poorly understood. We studied the sources of heterogeneity in the responses of human epithelial cells to oncogenic BRAF^V600E^ expression. We found that a narrow expression range of BRAF^V600E^ generated a wide range of activities of its downstream effector ERK. In population-level and single cell assays, ERK activity displayed a non-monotonic relationship to proliferation, with intermediate ERK activities leading to maximal proliferation. We profiled gene expression across a range of ERK activities over time and characterized four distinct ERK response classes, which we propose act in concert to generate the unique ERK-proliferation response. Altogether, our studies mapped the input-output relationships between ERK activity and proliferation providing important insights into how heterogeneity can be generated during OIS.

## INTRODUCTION

Activation or aberrant regulation of oncogenes promotes cellular transformation and tumorigenesis, enabling cancer cells to grow and avoid programmed cell death (Hanahan and Weinberg, 2011). For instance, activating RAS or RAF mutations (found in ~27% and ~8% in all human cancers, respectively) bypass requirements for growth factors, resulting in constitutive mitogenic signaling through the MAPK pathway (Hobbs et al., 2016; Holderfield et al., 2014). However, when proteins carrying oncogenic mutations are expressed ectopically in non-transformed cells, they cause the cells to enter a state of stable cell cycle arrest, a phenomenon known as oncogene-induced senescence (OIS) (Collado and Serrano, 2010; Serrano et al., 1997). OIS, first reported in primary diploid fibroblasts with HRasG12V expression (Serrano et al., 1997), was later found to be caused by various oncogenes and reported in multiple *in vitro* cell systems as well as at the organismal level (Collado and Serrano, 2010). Moreover, in mouse models, low-level expression of HRasG12V drives hyper-proliferation whereas high-level expression drives cellular senescence (Sarkisian et al., 2007). In cultured melanocytes, expression of BRAF^V600E^ initially stimulated moderate proliferation (3-7 days), which was followed by a progressive decrease in growth rate and eventual cell cycle arrest (Michaloglou et al., 2005). The cell cycle arrest typically involves the p53/p21WAF1 and p16INK4A/RB tumor suppressor genes and their interacting networks, although the roles of these proteins appears to be cell type- and context-dependent (Adams, 2009; Mooi and Peeper, 2006). OIS is currently considered to represent a bona fide tumor suppressor mechanism, acting alongside cell death programs.

Cell-to-cell heterogeneity is often observed during OIS, with some cells in a culture arresting and others continuing to proliferate. *In vivo*, malignant tumors are found adjacent to benign tumors despite the presence of the same driver oncogene mutation in both (Adashek et al., 2020; Michaloglou et al., 2008). In these cases, senescence-associated markers are only found in the benign or premalignant lesions and are progressively lost as the lesions become malignant. At a population level, the time between oncogene expression and OIS varies from a few days to several weeks and proceeds asynchronously (Michaloglou et al., 2005; Serrano et al., 1997). It remains unclear why a subset of cells in a population is better able to tolerate the negative effects of oncogene activation. Contributing factors are likely to include the type, strength and duration of the senescence-inducing signal, non-cell autonomous influences from oncogene expression, and the susceptibility of a cell to (epi)genetic reprogramming (Adams, 2009; Ruiz-Vega et al., 2020).

Studying how heterogeneity arises in OIS is important for understanding the initiation of cancer and devising effective therapies. In the current study, we focus on OIS induced by BRAF^V600E^, an oncogenic variant of a MAPK serine/threonine kinase that lies immediately upstream of MEK and ERK. MAPK activity induces the expression of multiple transcription factors that promote expression of positive regulators of the cell cycle, leading to cell cycle entry (Meloche and Pouysségur, 2007). BRAF^V600E^ is found in ~60% of cutaneous melanomas and is the primary target for the current standard of care for treating this disease. However, the same mutation is found in ~90% of melanocytic nevi, the benign, pigmented ‘moles’ found on the skin of most individuals. Thus, although activation of the MAPK cascades is a critical step in initiating melanocytic neoplasia it is not sufficient, perhaps because the induction of OIS prevents tumor formation (**Fig 1A**) (Davies et al., 2002; Pollock et al., 2003).

**Figure 1.**
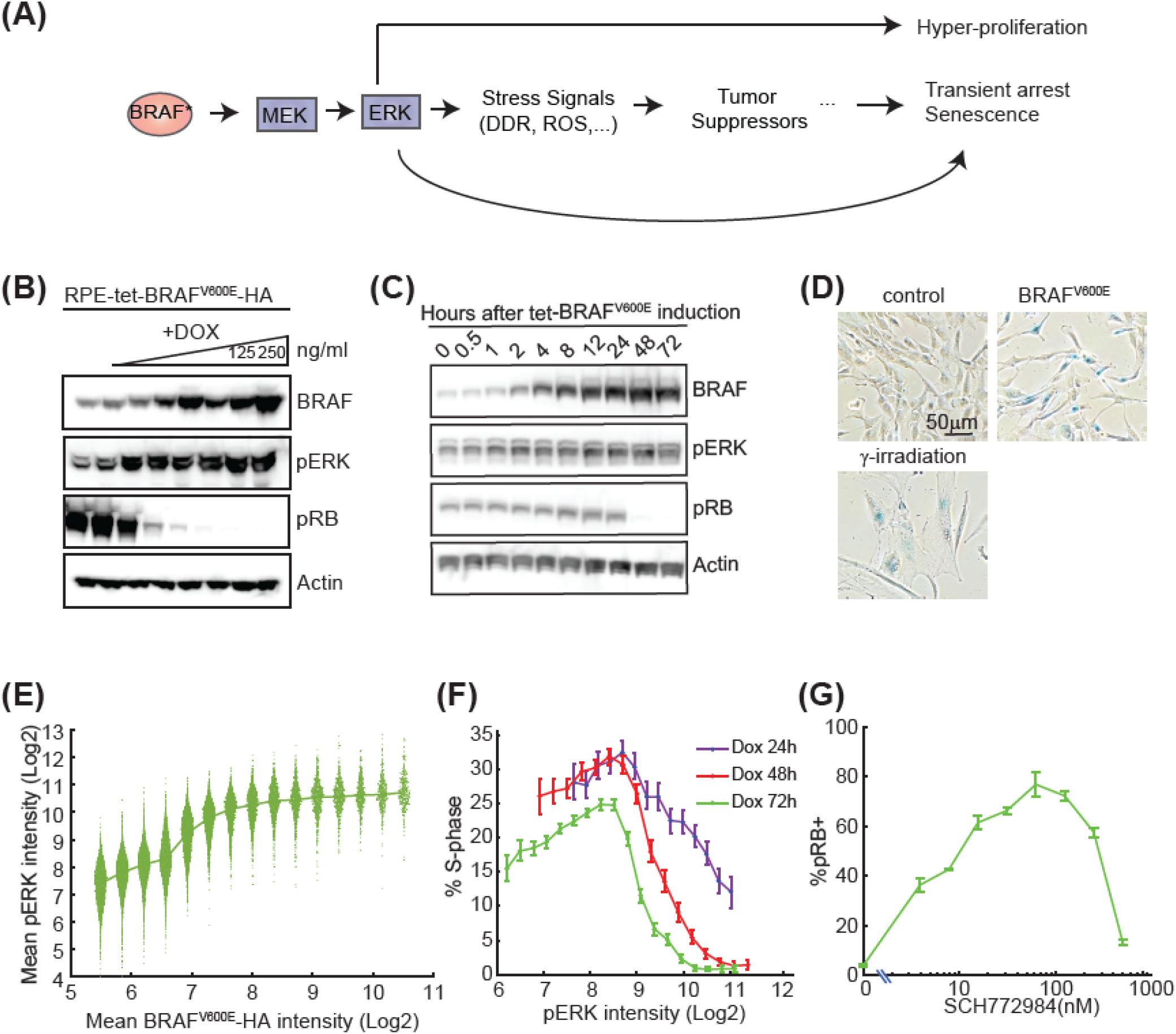
BRAF^V600E^-ERK pathway activation results in a non-monotonic proliferation response. (A) Schematics of pathways activated by oncogenic BRAF (* denotes oncogenic mutations in BRAF including V600E in this study). DDR, DNA damage response. ROS, reactive oxygen species. (B) Western blot analysis of levels of BRAF, active pERK, and the proliferation marker pRB following BRAF^V600E^ induction by increasing doses of doxycycline (DOX, 0-250ng/ml, 2-fold dilution from the right) for 72h. Actin is shown as a loading control. (C) Western blot analysis of levels of BRAF, active pERK, and the proliferation marker pRB at the indicated time points following BRAF^V600E^ induction with 250ng/ml DOX. Actin is shown as a loading control. (D) Representative images of cells assayed for senescence associated □-galactosidase (SA-□-gal) activity 7 days after DOX induction of BRAF^V600E^-HA at 250ng/ml or after 10 Gy □-irradiation. (E) RPE/tet-BRAF^V600E^-HA cells were treated with serial doses of DOX (0-250ng/ml, 2-fold dilution, also see **Fig S1A**) for 72h and immunostained for BRAF^V600E^-HA and pERK. The immunofluorescence data from all DOX doses were pooled together and single-cell pERK levels were extracted for equally spaced bins of BRAF^V600E^ expression. Each bin contains at least 400 cells. Each dot represents a single cell. (F) RPE/tet-BRAF^V600E^-HA cells were treated with serial doses of DOX as in (E) for 24h, 48h or 72h before immunostaining. The immunofluorescence data from all DOX doses were pooled together and the percent of cells in S phase (%S) was calculated for equally spaced bins of ERK activity (mean ± 95% bootstrap confidence interval). (G) RPE/tet-BRAF^V600E^-HA cells were treated with DOX (250ng/ml) together with different doses of ERK inhibitor (ERKi) (SCH772984, 0-500nM, 2-fold dilution) for 72h and then stained for pRB as a marker of proliferation. The percentage of pRB+ cells at each dose of ERKi is shown (mean ± SD of three replicate wells).

Consistent with this idea, expression of BRAF^V600E^ has been reported to induce senescence in a variety of cell lines including fibroblasts and melanocytes (Michaloglou et al., 2005; Vizioli et al., 2011; Zhu et al., 1998). ERK, the downstream effector of BRAF, plays a central role in proliferation decisions. Hyperactivation of the ERK pathway causes accumulation of cyclin-dependent inhibitors, but the overall input-output relationship between ERK activity and proliferation has not been established (Deschênes-Simard et al., 2014; Meloche and Pouysségur, 2007).

In this paper we investigated the relationship between BRAF^V600E^ levels, ERK activity and cell proliferation in non-transformed human hTERT-immortalized retinal pigment epithelial (RPE) cells. We showed that a narrow expression range of BRAF^V600E^ protein generated a wide range of ERK activities. We further showed a non-monotonic relationship between ERK activity level and proliferation response. We examined the source of the non-monotonic response through global transcriptional profiling, which revealed four categories of cellular responses across different ranges of ERK activities. Our study highlights the various networks of genes that are induced in response to differential ERK signaling, and provides important clues as to how individual or combinatorial classes of genes can generate a biphasic proliferation response.

## RESULTS

### The relationship between ERK activity and proliferation is non-monotonic

To establish a model of oncogene-induced senescence, we expressed the oncogenic BRAF^V600E^ variant carrying a C-terminal HA tag in RPE cells under the control of a doxycycline (DOX)-inducible promoter (**Fig 1B)**. Activation of the MAPK cascade by BRAF^V600E^ was assessed by measuring phospho-ERK (pERK) levels using western blotting. Time course analysis showed that, at a saturating dose of DOX (250ng/ml), BRAF^V600E^ levels increased for the first 24 hr and then plateaued, while pERK levels plateaued around 2 hr, and pRB (a marker of cell cycle progression) was undetectable by 48 hr (**Fig 1C**). To determine if BRAF^V600E^ expression induced senescence, cells were stained for *β*-galactosidase. Cells exposed to *γ*-irradiation, a well-established inducer of senescence, were used as a positive control. BRAF^V600E^ induction for 7 days resulting in *β*-galactosidase expression in cells comparable to irradiated cells, and higher than untreated controls (**Fig 1D**). These results suggest BRAF^V600E^ expression in RPE cells causes cell cycle exit and promotes senescence, consistent with induction of OIS.

We next investigated the relationships between BRAF^V600E^ expression, ERK activity, and proliferation outcomes. To induce variable BRAF^V600E^-HA expression levels, RPE/tet-BRAF^V600E^ cells were treated with various doses of DOX for 72 hr and then stained for HA, pERK, and EdU incorporation (a marker for DNA synthesis) (**Fig S1A-C**). Consistent with **Fig 1B**, proliferation was inhibited by BRAF^V600E^ expression in a DOX dose-dependent manner, with maximum proliferation occurring in the uninduced condition (**Fig S1C**). The data were then pooled for subsequent analysis of ERK activity, in which we binned BRAF^V600E^ expression levels measured in single cells (irrespective of the DOX dose) and quantified the ERK activity in each bin. This analysis revealed that pERK levels increased with BRAF^V600E^ expression at lower levels but then plateaued at higher levels, suggesting that ERK activity is saturated at intermediate levels of BRAFV600E overexpression (**Fig 1E**). Moreover, for any given level of BRAF^V600E^, pERK levels varied, demonstrating substantial cell-to-cell variability in activation of the MAPK cascade (**Fig 1E**). To determine the relationship between pERK levels and cell cycle progression, DOX was added to the RPE/tet-BRAF^V600E^ cells at various doses for 24, 48 or 72 hours and the fraction of S-phase cells was then quantified by EdU incorporation (**Fig 1F**). Binning the data on pERK levels (regardless of DOX dose), revealed that the fraction of cycling cells was highest at intermediate pERK levels, and decreased at higher and lower pERK values. This result suggests that a moderate induction of pERK enhances proliferation; however, beyond a certain level, proliferation is inhibited (**Fig 1F**). In the absence of BRAF^V600E^ expression, ERK activity levels fell in the lower range (between 6-9 Log2 intensity), and the fraction of proliferating cells associated with these levels ranged from 15-25% (**Fig S1B** & **S1D**). After BRAF^V600E^ induction, ERK activity levels fell into a higher range (between 8.5-11.5 Log2 intensity), and the associated proliferation frequencies ranged from 25% at the lower end of ERK activity to 1% at the higher end (**Fig S1D**). Because proliferation is highly sensitive to ERK activity in the range experienced by cells expressing BRAF^V600E^, it is likely that the heterogeneity in ERK activity during OIS (**Fig S1B**) leads to heterogeneous responses in terms of proliferation and senescence. To further confirm the non-monotonic relationship between ERK activity and cell proliferation, we over-expressed BRAF^V600E^ at a level sufficient to arrest most cells, and treated them with a dose series of ERK inhibitor (ERKi) (SCH772984) to titrate down ERK activity. Consistent with **Fig 1F**, the fraction of cycling cells (indicated by the pRB^+^ fraction) was highest at an intermediate ERKi concentration. By contrast, the fraction of proliferating cells was reduced when cells were treated with higher or lower ERKi concentrations (**Fig 1G**), suggesting a non-monotonic relationship between proliferation and ERK activity.

### Establishment of a new cell cycle reporter that differentiates G1, S, and G2 phases

Our data suggest that cells can make proliferation or arrest decisions in response to ERK activity. The data presented in **Fig 1** show the average response of a population of cells. We hypothesized that at the single cell level, internal cellular states such as cell cycle phases or the level or dynamics of oncogenic signaling, may influence the proliferation status. We sought to monitor ERK activity and its relationship to cell cycle progression using live-cell imaging. Commonly used live-cell cell cycle reporters, such as Geminin, monitor the G1-S transition (Sakaue-Sawano et al., 2008) but growing evidence suggests that G2 plays a pivotal role in proliferation-quiescence decisions (Min et al., 2020; Spencer et al., 2013; Yang et al., 2017).

To distinguish G1, S and G2 cell cycle phases we developed a biosensor based on the PCNA-interacting-protein (PIP)-box motif. PIP boxes are recognized by Cul4^Cdt2^ E3 ubiquitin ligase and are degraded specifically at S-phase (**Fig 2A**) (Havens and Walter, 2009). Because ectopic expression of human PIP-box containing proteins has the potential to interfere with normal cell cycle progression, we developed a sensor based on the PIP-box motif of *Drosophila* E2F (dE2F) (Shibutani et al., 2008). The sensor includes the N-terminus of dE2F (amino acids 1-187) fused to the red fluorescent protein (FP) mCherry. The resulting mCherry-PIP protein contains PIP boxes functional in humans as well as a naturally occurring nuclear localization signal. RPE cells stably expressing mCherry-PIP, a Turquoise FP-tagged nuclear histone marker (H2B-Turq), and a Venus FP-tagged Geminin (1-110) reporter were established. The Geminin (1-110) reporter accumulates in S-phase and is degraded in G1 phase (Sakaue-Sawano et al., 2008). mCherry-PIP exhibited periodic accumulation in the nucleus throughout 48 hr of imaging (**Fig 2B)**.

**Figure 2.**
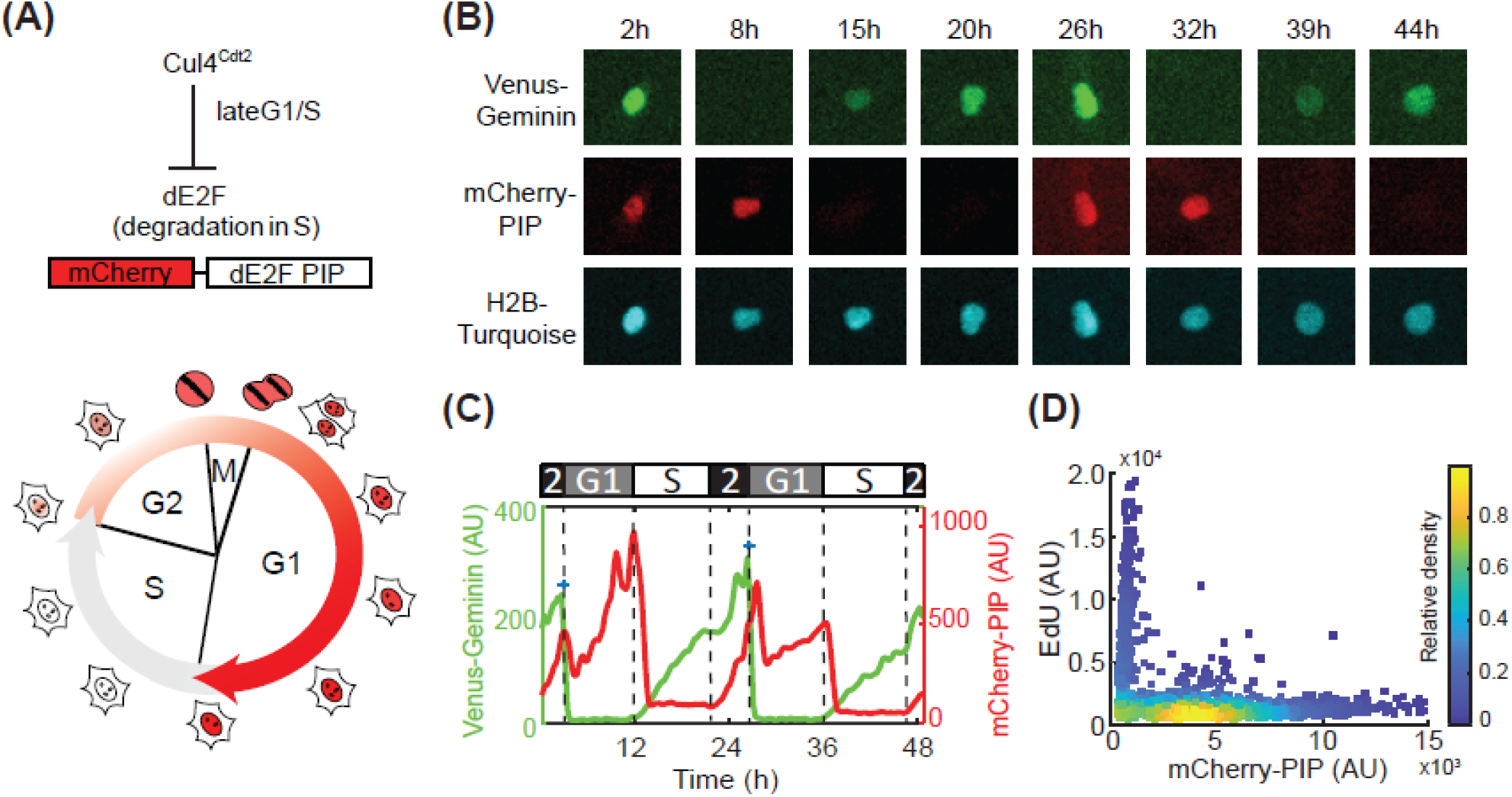
Development and characterization of a live-cell sensor that identifies G1, S, and G2 cell cycle phases. (A) Schematic of PIP (PCNA-interacting protein) motif based S-phase biosensor. *Drosophila* E2F1 undergoes S-phase specific degradation by the highly conserved PIP motif in E2F1 and the Cul4 (Cdt2) E3 ligase (top). The bottom diagram shows cell cycle progression with nuclear mCherry-dE2F PIP fluorescence changes. (B) Images of an RPE cell stably expressing Venus-Geminin (1-110), mCherry-dE2F PIP and H2B-Turquoise proceeding through the cell cycle during the 48 hr imaging time. (C) Single cell trace of mCherry-dE2F PIP (red) and Venus-Geminin (1-110) (green) shown in (B). (D) RPE cells stably expressing mCherry-dE2F PIP were pulsed with EdU for 15min before fixation and detection of mCherry-dE2F PIP and EdU. Density scatter plots show mCherry-dE2F PIP intensity versus EdU fluorescence intensity. Each dot represents a single cell. The color scale represents the relative density.

When comparing both cell cycle reporter levels within the same cell, mCherry-PIP rapidly dropped in expression when Venus-Geminin (1-110) began to accumulate at the G1-S transition (**Fig 2B** & **C**). The mCherry-PIP fluorescence signal rose subsequently while Venus-Geminin continued to accumulate. Thus, the patterns of Venus-Geminin (1-110) and mCherry-PIP protein accumulation and degradation are consistent with the anticipated properties of the reporter proteins (Havens and Walter, 2009; Sakaue-Sawano et al., 2008). By labeling S-phase cells with EdU, we validated that mCherry-PIP proteins were lowest in S-phase (EdU positive) and were only present when cells were not in S-phase (**Fig 2D**). Thus, in live-cell experiments G2 can be identified by the presence of both Venus-Geminin (1-110) and mCherry-PIP (**Fig 2C**). In addition, cell cycle phases can be computationally derived from live-cell data by quantifying only the levels of the mCherry-PIP reporter as follows: G1 corresponds to the period between nuclear division and a rapid drop in mCherry-PIP fluorescence; S corresponds to the period between this rapid drop and right before resynthesis of mCherry-PIP occurs; and G2 is corresponds to the period in which mCherry-PIP accumulates prior to the next cell division (see material and methods for details of this analysis; **Fig 2C**).

### Live imaging traces revealed a bell-shaped relationship between ERK activity and cell cycle entry in single cells

Given the presence of cell-to-cell heterogeneity in ERK levels and OIS induction, we sought to establish the relationship between ERK activity, cell cycle phase transitions, and cell fate at a single cell level. We therefore generated an RPE cell line stably expressing DOX-inducible BRAF^V600E^-HA, mCherry-PIP, and EKAREN5, a reporter for ERK activity. This line (which we termed BRAF^V600E^ Dual Reporter cell line) allowed us to induce oncogenic BRAF^V600E^ and simultaneously measure ERK activity and cell cycle progression in the same cell through long-term live imaging. EKAREN5 (Ponsioen et al., 2021) is a version of the widely used EKAREV FRET-based ERK activity reporter (Komatsu et al., 2011; Ponsioen et al., 2021) that has been engineered to make it insensitive to CDK1/cyclin B activity at G2 and M phases. Control experiments confirmed that EKAREN5 sensor reflects ERK activity in our RPE line and that activation during G2/M phase is strongly reduced relative to EKAREV (**Fig S2A)**.

We imaged asynchronous cultures of BRAF^V600E^ Dual Reporter cells for 24h to obtain cell cycle phase information under unperturbed conditions and then added DOX to induce BRAF^V600E^ expression, followed by live cell imaging for 3 days to monitor ERK activity and cell cycle progression. We used a semi-automated tracking method to label the timing of individual division events, and computationally derived the cell cycle phases and ERK activities (**Fig 3A;** see materials and methods). When BRAF^V600E^ Dual Reporter cells were treated with DOX (24h post imaging), a rapid increase in ERK activity was observed within 2-4 hours (**Fig 3B**, consistent with data in **Fig 1C**) while control cells showed basal ERK activity with occasional pulses throughout the imaging period (Gerosa et al., 2020). Most DOX-treated cells underwent one or two divisions prior to entering prolonged G1 arrest, whereas cells not treated with DOX continued to proliferate, serving as a control for the effects of long-duration imaging (**Fig 3C**). A detailed cell cycle duration analysis revealed that prior to the prolonged G1 arrest, the G1 and S phase lengths of the preceding cell cycle remained unaltered while G2 lengths (orange blocks in **Fig 3B**) increased (**Fig S2B**). The increase in G2 phase length was ERK-dependent, as addition of ERK inhibitor shortened G2 duration (**Fig S2C**).

**Figure 3.**
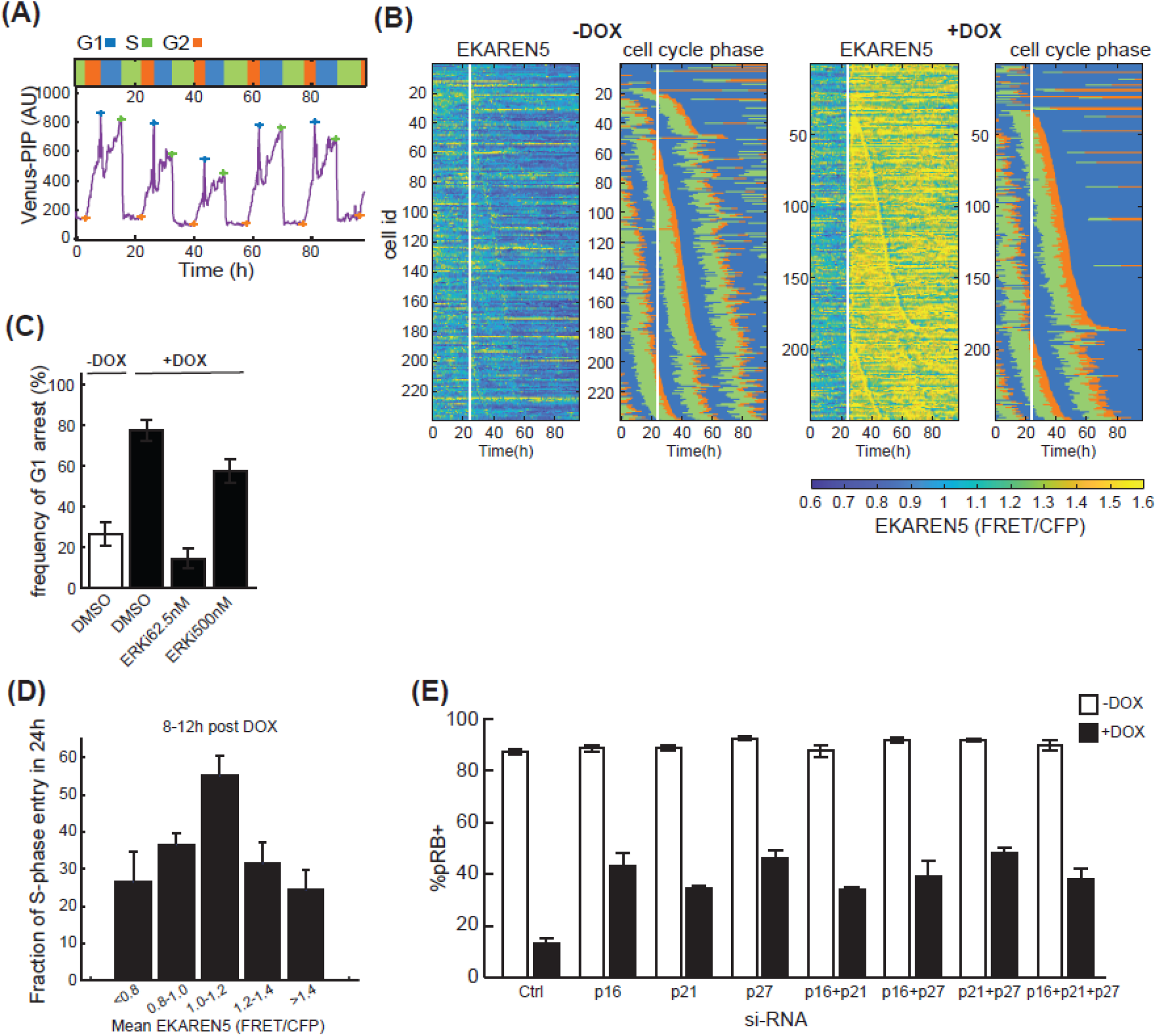
Single-cell live imaging traces revealed a bell-shaped correlation between ERK activity and proliferation. (A) Sample single cell trace of Venus-dE2F PIP in RPE cells proceeding through the cell cycle. Blue, green and orange crosses mark the start of G1, S and G2 phase. Respective length of each cell cycle phase is shown above. (B) Heatmaps of ERK activity (EKAREN5) and cell cycle distribution (the same color scheme as shown in (A)) in RPE reporter cells treated with control vehicle (-DOX, left) or DOX (+DOX, right). RPE Dual Reporter cells stably expressing tet-regulated HA-tagged BRAF^V600E^, ERK reporter EKAREN5 and mCherry-dE2F PIP were treated as indicated 24h after the start of imaging (white vertical line). Each horizontal line represents a single cell. (C) Frequency of G1 arrest in RPE Dual Reporter cells treated with control vehicle (-DOX) or DOX together with DMSO or ERKi. Cells and treatment schemes were the same as in (B) and the percent of G1-arresting cells was calculated following time-lapse imaging (mean ± 95% confidence interval, n>200 cells per condition). (D) Fraction of S-phase entry in response to increasing ERK activity. The RPE Dual Reporter cells in (B) were treated with ERK inhibitors (0, 62.5, or 500nM) in the absence or presence of DOX and at 24h after the start of live imaging. Cells were then imaged for another 72h. To broaden the sampling of ERK activity range, data from all treatments were pooled together and the mean ERK activity between 8-12 hr post treatment was calculated. The probability of entering into S-phase was quantified within 24 hr after the ERK span (mean ± 95% confidence interval, n>100 for each ERK activity bin). (E) Percentage of pRB positive cells after siRNA-mediated depletion of the indicated CDK inhibitors (individually or in combination) 2 days post ±DOX treatment in RPE cells stably expressing tet-HA-tagged BRAF^V600E^ (mean ± SD, n=4 replicates).

To map the relationship between ERK activity and cell cycle progression, cells were treated with ERKi at different doses at the same time as BRAF^V600E^ induction with DOX. While 77% of cells treated with DOX in the absence of ERKi underwent G1 cell cycle arrest with decreased total division numbers, the addition of 62.5nM ERKi rescued the arrest (**Fig 3C** & **Fig S2D**). However, at 500nM ERKi, the fraction of cells undergoing G1 arrest increased, consistent with a non-monotonic relationship between ERK levels and proliferation (**Fig 3C** & **Fig S2D**). To quantify this relationship in single cells, we pooled single-cell trajectories based on mean ERK activity, and then computed the fraction of cells that entered S-phase within the following 24 hr window. Mean ERK activity was determined between 8-12 hr post BRAF^V600E^ induction when ERK activity levels stabilizes in cells (**Fig 1C** & **Fig S3A**). When the probability of S-phase entry was plotted against mean ERK activity, we again observed a nonmonotonic, bell-shaped response curve (**Fig 3D**). This relationship held for the subsequent time intervals measured between 12-16 hr and 16-24 hr post DOX addition (**Fig S2E**). The high sensitivity of OIS to increases in ERK activity above the optimum value likely explain cell-to-cell heterogeneity in response to BRAF^V600E^ overexpression.

Previous OIS studies have suggested that activation of p16INK4A and p53 are the two major mechanisms leading to cell cycle arrest (Adams, 2009; Mooi and Peeper, 2006). To investigate this possibility in RPE/tet-BRAF^V600E^ cells, we used RNAi to acutely knock-down p16, p21 (a downstream target of p53) or p27 alone or in various combinations. The CDK inhibitor p27 was included due to its well-documented role in integrating diverse signals that regulate cell-cycle exit (Chu et al., 2008). Cells were then treated with DOX and the fraction of cycling cells was measured (**Fig 3E**). We found that knockdown of CDK inhibitors individually and in combination had a modest but reproducible effect on BRAF^V600E^-mediated arrest, but in no case was proliferation full restored to control levels. These results imply that additional proteins, beyond those suggested by previous studies, are involved in OIS, prompting us to apply a more systematic approach.

### Deep RNA sequencing identifies genes that respond to ERK activity levels

We hypothesized that factors mediating ERK activity-dependent cell fate decisions must themselves undergo changes in expression or activity in response to varying levels of ERK activity. To systematically identify genes whose expression changes with ERK activity, we performed deep RNA sequencing of RPE/tet-BRAF^V600E^ cells treated with a combination of DOX for varying times (including 0 – an untreated control, and 1, 2, 4, 8, 16 and 24 hr) and ERKi at different concentrations (**Fig 4A**). The ERKi concentrations were chosen to sample the full range of proliferation responses based on previous titration experiments (**Fig 1G**). The time points were selected based on the observation that RPE cells showed a bell-shaped relationship between ERK activity and proliferation (**Fig 1F**) as early as 24h after BRAF^V600E^ induction. The early time points allow identification of genes that are more directly responsive to ERK activity changes, while the later time points reveal long-term effects. Of note, the live imaging experiments using BRAF^V600E^ Dual Reporter cells show that ERK activities peak 1-2 hr following DOX and ERKi treatment and then slowly decay while remaining at distinct levels during the subsequent 24h period of our experiments (**Fig S3A**). These results suggest our treatments resulted in fast and stable ERK responses. To evaluate the effects of ERK inhibition on normal cycling cells, cells were treated with different doses of ERKi for 24 hr without the induction of BRAF^V600E^. The resulting gene expression dataset involved 43 conditions assayed in duplicate. We detected ~13,000-14,000 coding transcripts in each condition (available in GEO: GSE180210) with an average Pearson correlation coefficient of 0.99 for replicates, demonstrating high reliability across the data (**Fig 4B** & **S3B**).

**Figure 4.**
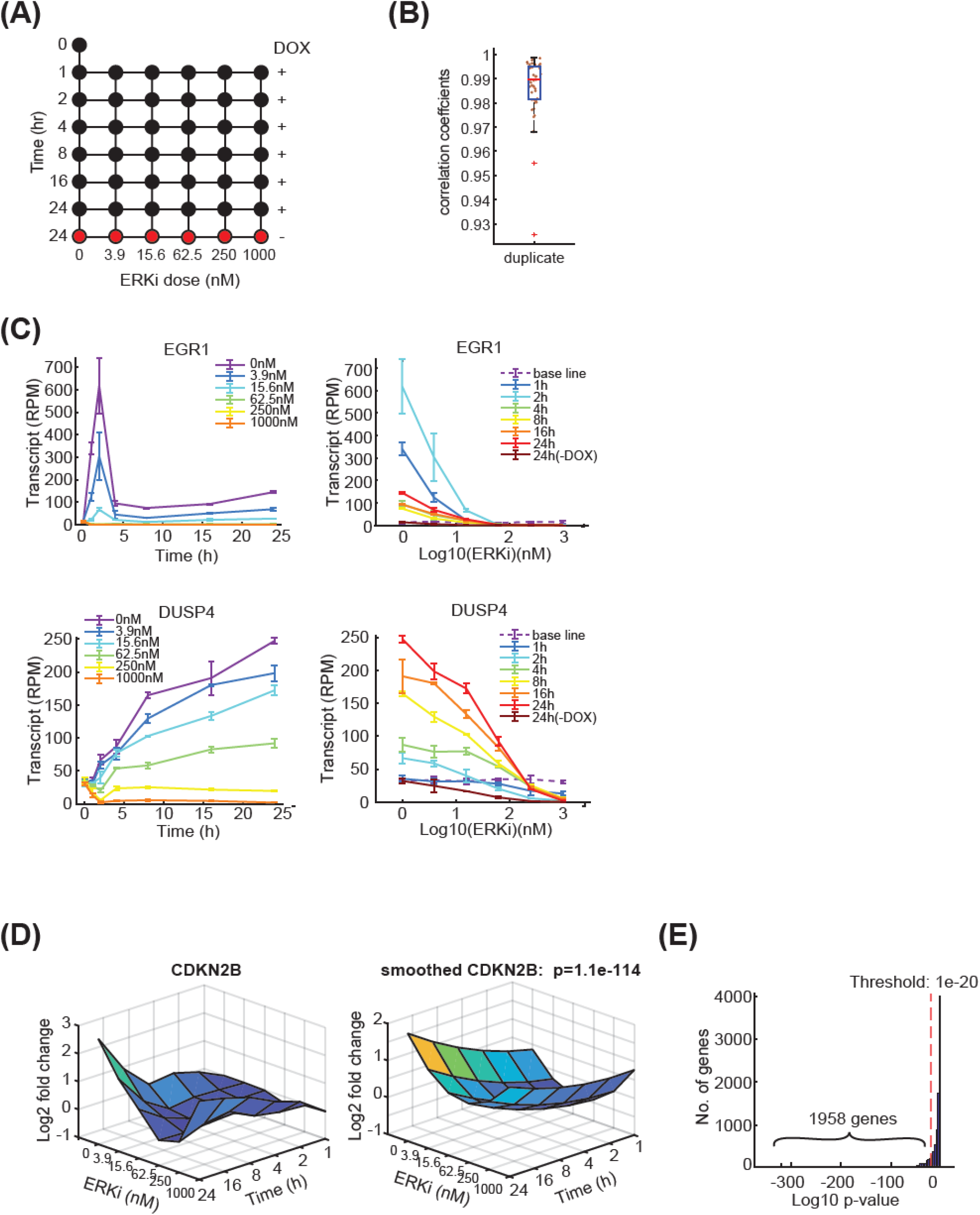
Deep RNA sequencing identifies genes with altered expression in response to different ERK activities. (A) RNA seq experimental design. RPE cells stably expressing tet-regulated HA-tagged BRAF^V600E^ were treated with DOX and variable concentrations of ERKi for 1-24h as indicated (43 conditions). Each condition was performed in two independent replicates (n=12 for no-treatment control). (B) Box plots showing correlation coefficients for RNA seq replicates. Each dot represents a pair of replicates. (C) Transcript levels as measured by RNA seq of EGR1 and DUSP4 in response to variable time of treatment (left) or variable ERKi concentrations (right) (mean ± SD from two independent replicates, base-line was plotted from no-treatment control). (D) (Left) Log2 fold change of CDKN2B transcripts as a function of treatment time and ERK inhibitor dose. Values were normalized to untreated control. (Right) Smoothed quadratic surface fit for CDKN2B expression. P-value shows the goodness of fit between the quadratic surface and a flat surface. (E) Selection of differential expressed genes based on quadratic surface fits as shown in (D). Genes with a p-value less than 1e-20, calculated as shown in (D), were considered differentially expressed.

As a first step in validating the approach, we quantified the levels of two genes, EGR1 and DUSP4, whose expression is known to be responsive to ERK activation (**Fig 4C**) (Amit et al., 2007). EGR1 exhibited a rapid and dramatic (~50-fold) induction within 2 hr of DOX treatment and then decreased rapidly, remaining at a level ~8-fold above its pre-induction levels for the duration of the experiment (**Fig 4C**, left). This time-course is consistent with its role as an immediate-early response gene. In contrast, DUSP4 rose steadily by ~6-fold over a 24 hr period, consistent with its role as an early-response gene mediating negative feedback in the MAPK signaling cascade (**Fig 4C**, left). Induction of EGR1 and DUSP4 fell in a dose-dependent manner when ERKi was present (**Fig 4C**, right). These results confirm that our transcript profiling studies have high dynamic range and can readily detect the induction of different ERK activity-dependent gene expression programs.

While it is common to analyze RNA-seq data to identify changes in expression associated with a single experimental variable (e.g. time or drug dose as in **Fig 4C**), given the dynamics of ERK activity, we sought to identify genes differentially expressed as a function of both time and ERKi dose. Comprehensive cross-correlations between different treatment conditions showed that samples collected at different times and/ or different ERKi doses can have similar transcriptional programs (high correlation), whereas samples collected at similar times and/ or ERKi doses can have low correlation in their transcriptional programs, exemplifying the complexity of our datasets (**Fig S3B**). Thus, inferring differential expression using traditional approaches poses a substantial challenge, since both doseresponse and temporal dynamics need to be accounted for. To address the challenge, we used regression with quadratic terms to identify the best-fitting time-dose response for every gene using QR factorization of the Vandermonde regressor matrix (Macon and Spitzbart, 1958) (**Fig 4D**). This type of factorization is an effective way to compute least-squares fits for a large number of genes, as computationally expensive factorization only needs to performed once for a specific set of dose-time combinations. Gene-specific regression then only required computationally cheap matrix-matrix multiplication. The purpose of computing the quadratic surface approximation is to minimize noise across the landscape of treatments and emphasize time and dose-dependent trends in the data.

To identify differentially expressed genes we compared the goodness of fit between the quadratic response surface for each gene and a flat surface. P-values were computed using standard likelihood ratio test and subjected to multiple-testing corrected using Bonferroni-Holm (Holm, 1979). Data for CDKN2B (the p15^INK4b^ kinase inhibitor) is shown in **Fig 4D** by way of illustrating the approach: CDNK2B expression is induced steadily over 24 hr and exhibits a U-shaped response to ERKi concentration, with a minimum expression at 62.5 nM (**Fig 4D**, left). Quadratic regression on the data yields a smoothed surface (**Fig 4D**, right) that is significantly different from a flat surface (p = 1.1e^-114^, likelihood ratio test). Using this approach, we identified 1958 genes that exhibited significant (p< 1e^-20^, likelihood ratio test) differential expression over time and ERKi dose (**Fig 4E**). GO analysis showed that these genes fall into different functional categories including extracellular matrix signaling, cancer pathways, DNA replication, and cell cycle control (**Fig S3C**). We investigated the presence of known BRAF^V600E^, ERK, and senescence signatures (**Fig 1**) in the 1958 differentially-induced genes using GSEA (Gene Set Enrichment Analysis) (**Fig S3D**). The differentially expressed genes identified by quadratic regression were significantly enriched for the senescence and MAPK signatures examined, validating the effectiveness of our analysis method.

### Characterization and clustering of gene expression in relation to ERK signaling dosage

To identify overall trends in the data, we performed principal component analysis (PCA) with the transcriptional data from all 43 conditions for the previously identified differentially expressed genes. The first two principal components (PC1 and PC2) explained 81% of overall variance in the data, with PC1 and PC2 capturing 54.7% and 26.5% of the variance respectively (**Fig 5A**). When plotting weights of PC1 and PC2 separately for every time point post treatment (**Fig 5A**), the variance of weights increased in a time-dependent manner, starting at 4 hr (top right panel) and progressively forming a bell-shape that reached its full extent at the 24 hr time point (bottom right panel). Interestingly, the bell-shaped curve at 24 hr closely resembled the phenotypic correlation between ERK activity and proliferation response (**Fig 1F**, **1G** & **Fig 3D**). Based on these findings PC1 appeared to correspond to differences in ERK activity and PC2 to differences in proliferative index.

**Figure 5.**
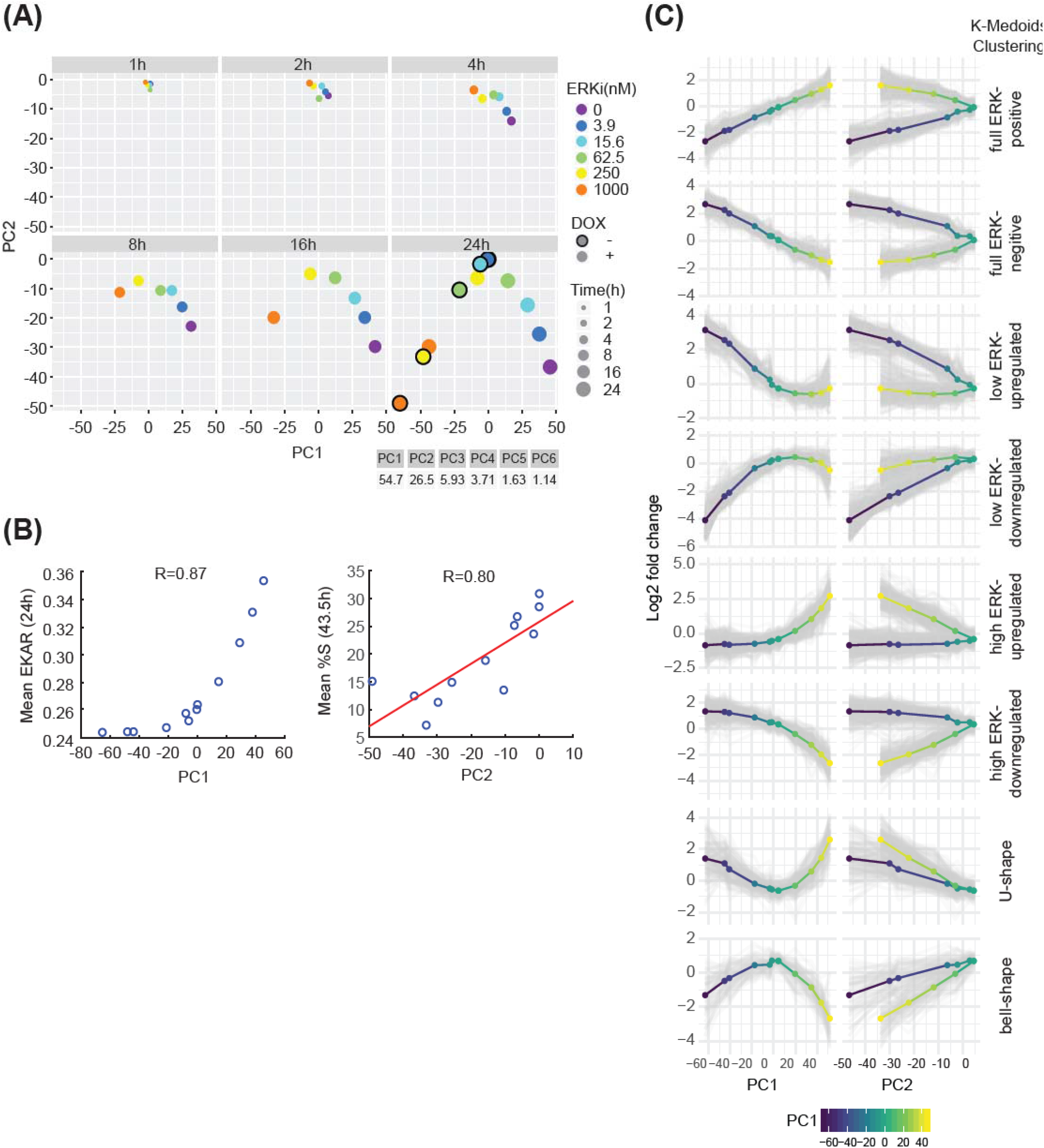
Gene classification by clustering of gene expression reveals different response types to changing levels of ERK activity and proliferation. (A) Principal-component analysis (PCA) of log2 fold-change in gene expression vs untreated control for each treatment (mean of two replicates) from RNA-seq. PC1 (principle component 1) and PC2 (principle component 2) contribute 54.7% and 26.5% of the variance in the data, respectively. (B) Live imaging experiments revealed the correlation between PC1 and ERK activity (EKAREN5) and the correlation between PC2 and proliferation (percent of cells in S-phase, %S). BRAF^V600E^ Dual Reporter cells were treated with ERKi (0, 3.9, 15.6, 62.5, 250, 1000nM) in the absence or presence of DOX at the start of live imaging and continued imaging for 43.5h (12 treatments). PC1 values obtained at 24h post treatments in (A) were plotted against the ERK activity measured at 24h with comparable treatments from live imaging experiments. Similarly, PC2 values at 24h post treatments were plotted against the percent of cells in S-phase measured at 43.5h from live imaging experiments. 43.5h was chosen to account for the time delay between gene expression and cell cycle entry. (C) K-medoids clustering of the 1958 differential expressed genes identified from RNA seq experiments. Log2 fold-change expression data of each differential expressed gene at 24h were clustered by k-medoids clustering (k=8). Each cluster has a characteristic gene expression pattern in response to varying levels of PC1(a proxy of ERK signaling) and PC2 (a proxy of proliferation index). Mean expression levels of each cluster are shown as multi-colored line (blue-yellow represents low-high PC1 values).

To test this hypothesis, we measured the mean ERK activity and degree of proliferation at each condition assayed by RNA-seq. We treated BRAF^V600E^ Dual Reporter cells with DOX and different doses of ERKi simultaneously (mirroring the experimental conditions in RNA seq). We then performed live-imaging experiments, and monitored mean ERK activity and fraction of cells in S-phase along the live imaging trajectories. PC1 values (**Fig 5A**, bottom right) were highly correlated (R=0.87) with mean ERK activities measured under the same conditions using the EKAREN5 reporter (**Fig 5B**, left). Moreover, the fraction of cells in S-phase (measured using mCherry-PIP) was highly correlated with the value of PC2 (R= 0.8; **Fig 5B**, right). Thus, we conclude that the first principal component is a proxy for ERK activity whereas the second principal component is a proxy for proliferation index. Moreover, we found that the value of PC2 was similar for conditions that had similar average ERK activity, regardless of how that activity level was achieved. For example, a PC2 value of approximately −30 was achieved in cells not expressing BRAF^V600E^ and treated with low dose (250 nM) ERKi, as well as in cells with BRAF induced and treated with high dose ERKi (1000 nM) (**Fig 5A**, 24h plot, yellow and orange circles). These data also strongly suggest that the two primary drivers of gene expression, over a wide range of conditions, are the ERK activity level and the extent of proliferation.

To investigate the possibility of non-specific ERKi targets amongst the 1958 differentially expressed genes, we determined whether the effects of the inhibitor on these genes could be rescued by boosting ERK expression. Of the 1958 genes, many were differentially expressed in the presence of 250nM ERKi (compared to untreated cells). However, gene expression was restored to resemble that in the control when cells were treated with 250 nM ERKi plus DOX to induce BRAF^V600E^ and ERK activity (**Fig S4A**). Furthermore, comparing two conditions with similar ERK activities (ERKi 250 nM -DOX vs ERKi 1000 nM +DOX) (**Fig S4B**, left) revealed a high correlation (R^2^ = 0.97, **Fig S4B**, right), suggesting that the set of differentially expressed genes was primarily responding to ERK activity. These results together suggest that ERKi effects can be rescued by overexpression of BRAF^V600E^ and that the potential off-target effects of ERKi are likely very minimal.

We expected expression levels of genes to have a differential patterned response to varying levels of ERK signaling (PC1) and proliferation rates (PC2). To further subdivide gene expression programs, we used unsupervised k-medoids clustering based on PC1 and PC2 values at the 24 hr time point (**Fig 5C**). K-medoids is a variant of k-means clustering which is robust to outliers and also allows the use of arbitrary distance metrics. We computed pairwise Euclidean distances for the log2 fold-changes (relative to an untreated control) in the set of 1958 genes and then performed k-medoids clustering. With k = 8 clusters, differences in relationship between changes in expression and PC1 or PC2 values were evident (**Fig 5C**). For instance, in the “full ERK-positive” cluster, the gene expression goes up with increasing PC1 value suggesting a strong positive correlation. Yet, in the same set of genes, a single PC2 value can have two different levels of gene expression (from treatments that have either high or low PC1 values) indicating a weak correlation between gene expression and PC2 value. Similar clustering results were obtained with data from all the time points.

The results of k-medoids clustering yielded clusters that could be grouped by visual inspection into four qualitatively different response classes to ERK activity, each having two clusters with opposite trends (**Fig 6**). Class I included genes whose expression is correlated either positively (n=283, 14.5%) or negatively (n=296, 15.1%) with ERK activity across its full range, resulting in a linear relationship (“ fullrange ERK responder”; **Fig 6A** red lines). Canonical negative feedback regulators of the MAPK pathway such as DUSP4/6 and SPRY2 belonged in this group (Amit et al., 2007). Class II genes were those in which differential gene expression fell with ERK activity in a non-linear, “convex” manner (blue lines). At low ERK activity levels, 248 (12.7%) were significantly upregulated and 404 (20.6%) were down regulated. Class II included genes involved in DNA replication, DNA damage repair and the G1/S cell cycle transition (e.g. G1 cyclins) (**Fig S5A**). Class III was similar to the Class II with the response window shifted to higher ERK activity ranges (green lines). This class included 281 upregulated genes (14.4%) and 231 (11.8%) down-regulated genes. Class III included genes involved in differentiation, migration/motility, cytokine response and growth factor activity. Class IV (orange lines) comprised genes with a bell-shaped (n=71, 3.6%) or U-shaped (n=144, 7.4%) response curves, having the greatest differential gene expression at the lowest and highest ERK levels. This class included the CDK inhibitor CDKN2B (p15). Expression of Class IV genes was highly dependent on PC2 as seen by a monophasic response where similar expression levels were obtained for a given PC2, regardless of PC1 value (**Fig 5C**, bottom). The classifications of gene expressions in **Fig 6A** exemplify different strategies cells utilize in response to varying levels of ERK activity.

**Figure 6.**
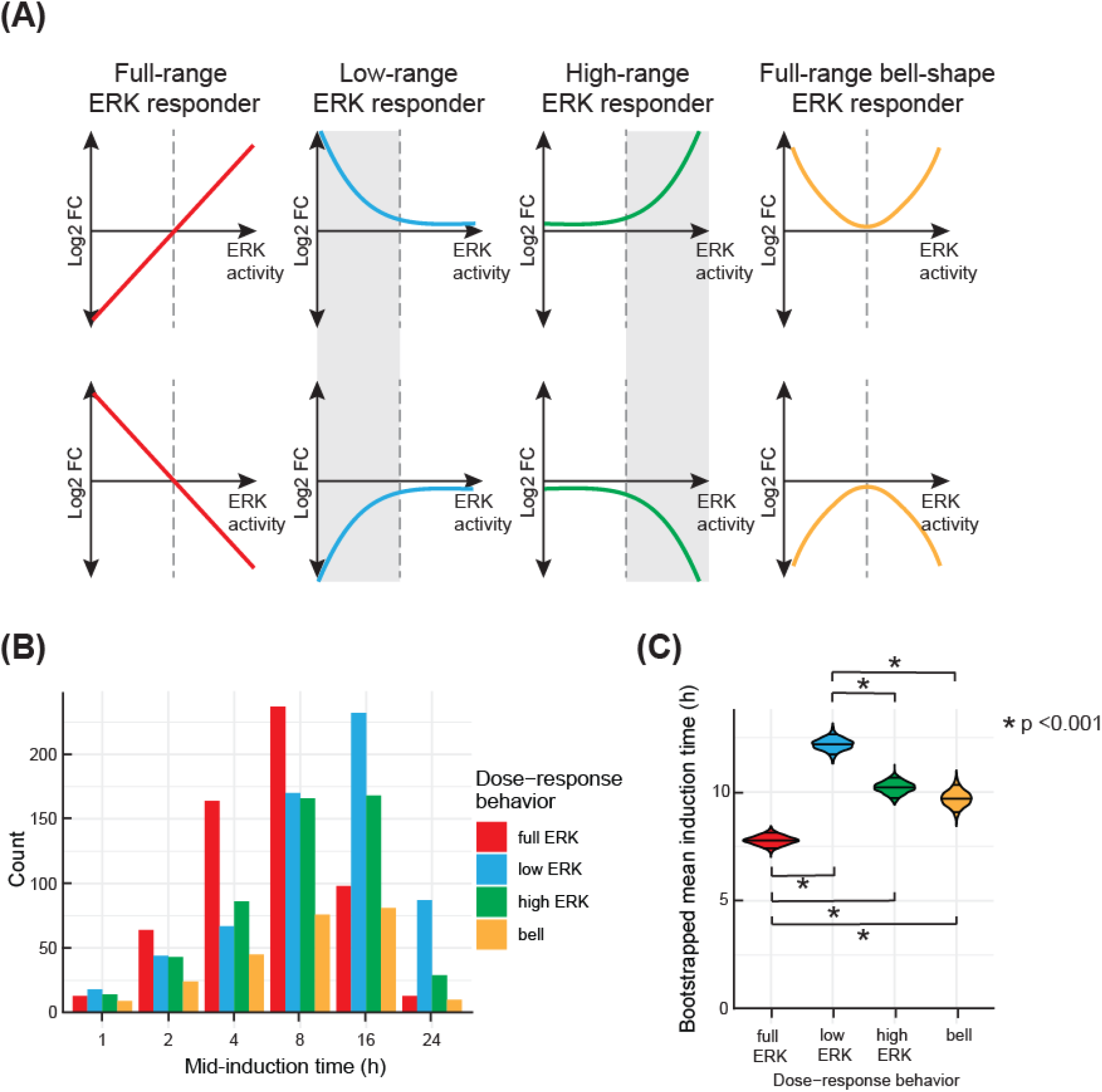
Patterns of ERK-dependent gene expression clustered by ERK activity and time of induction. (A) Four distinct classes of gene expression patterns in response to variable ERK activity. Fullrange ERK responder class includes genes with gene expression proportional to ERK activity across the entire range of activity levels. Low-range ERK responder class includes genes that up- or down-regulate their expression at lower range of ERK activity but remain non-responsive at higher ERK activity. High-range ERK responder class shows differential gene expression only at higher ERK activity levels. Full-range bell-shape ERK responder class includes genes that have a bell or a U-shaped response with the most extreme responses happening at low and high ERK activity. For illustration purposes, we substituted ERK activity for PC1 value since the two are highly correlated. (B) Temporal dynamics of each class of genes shown in (A) in response to DOX/ERKi treatment. Genes in each class were grouped according to the earliest time-point at which they achieved 50% of the maximal change observed over the course of the experiment (see Materials and Methods). The time-point of mid-induction is shown on the x-axis and the number of genes falling into each category is shown on the y-axis. (C) Mean induction time of ERK responder classes. The violin plots show bootstrapped estimates of the mean for each cluster. Significant differences in the induction time between ERK-response clusters are highlighted with black bars. P-values are derived from ANOVA followed by Tukey’s honest significant difference test.

We also investigated the relationship between gene expression profiles and time. To obtain a simple measure of the timing of gene induction or repression, we considered the time at which each gene first reached half of its maximal differential gene expression value in a weighted average that aggregated data across different ERKi doses (see Materials and Methods). Genes were grouped according to the time of this absolute half-maximum change in expression (the “mid-induction time”). This analysis revealed genes responding on rapid (1-2 hr), intermediate (4-8 hr) and slow (16-24 hr) time scales (**Fig S5B**). For instance, the previously mentioned EGR1 and DUSP4 exhibit rapid and intermediate time scales of induction, respectively (Amit et al., 2007). In general, genes in any ERK response category from **Fig 6** were found in any temporal response category. However, we found that genes in the fullrange ERK responder group (Class I; red lines in **Fig 6A**) tended to respond at earlier times in BRAF^V600E^ induction than genes in Class III or Class IV. Genes in the low-ERK responder group (Class II) responded most slowly to BRAF^V600E^ induction (**Fig 6C**). These results suggest only modest correlations between time- and ERK activity-dependent gene regulation.

We envision that combinations of different temporal and ERK dose-response classes contribute to the overall bell-shaped proliferation response. For instance, low G1 cyclin expression (low ERK down-regulated group) combined with high CDK inhibitor expression (U-shape group) at low ERK activity could trigger cell cycle arrest. Similarly, upregulation of CDK inhibitor at high ERK activity (U-shaped group) could also trigger cell cycle arrest. Even though the proliferation response is stable over time (**Fig 1F**), the underlying mechanisms could vary over time, as we detected a large variation in the kinetics of ERK-responsive genes. Thus, a combination of ERK activity-responsive genes could function collaboratively to achieve heterogeneous proliferation responses within a population of cells, across time or ERK activity levels.

## DISCUSSION

While OIS is an efficient way to halt tumor development, it is not uncommon to see genetically homogenous cells respond to oncogenes in an asynchronous and heterogeneous manner. In this paper, we showed that cell to cell variability in OIS induction can be traced back to differences in ERK activity at a single cell level. A narrow range of ectopic BRAF^V600E^ expression generates cells with a wide range of ERK activities. Such “noise” is common in intracellular signaling networks (Suderman et al., 2017) and it is well established that cell-to-cell differences in protein levels can arise from transcriptional bursting and unequal partitioning of cytoplasm at cell division. These generate a lognormal distribution of protein concentrations across a population of cells that is sufficient to cause dramatic differences in their downstream protein activity and cell fate (Spencer et al., 2009). Using a newly developed single-cell PIP reporter and live-cell imaging, we found that entry into senescence is preceded by a prolonged G2 phase but not by a burst of proliferation as previously described (Courtois-Cox et al., 2008; Ogrunc et al., 2014). Single cell imaging revealed a non-monotonic, bell-shaped relationship between ERK activity and cell proliferation. Our data showed that proliferation is highly sensitive across the entire range of ERK activity, and thus, a population of cells expressing BRAF^V600E^ will continuously exhibit differences in ERK activity and heterogenous outcomes with some cells continuing to proliferate and others undergoing OIS. We speculate that cells that can proliferate in the presence of elevated BRAF^V600E^ are likely to give rise to tumors. It is also possible that cells that are induced to arrest in the presence of high ERK activity could once again become proliferative in the presence of the BRAF and MEK inhibitors used therapeutically.

Our data support a Goldilocks principle (“just the right amount”) for ERK activity such that an intermediate amount promotes proliferation, whereas much higher or lower levels prevent it. Accordingly, hyperactivating mutations in ERK are much less common than those in RAS and RAF (Deschênes-Simard et al., 2014; Roberts and Der, 2007), as ERK mutations are likely to lead to OIS. On the contrary, hyperactivation of its upstream regulators, e.g. RAS and RAF, are commonly found as oncogenic drivers (Hobbs et al., 2016; Holderfield et al., 2014), suggesting they may promote complex gene expression programs that either limit ERK activity or bypass the arrest caused by high ERK levels. Based on the Goldilocks principle, one can also predict that increasing ERK activity in cancer cells will likely push cells into senescence or cause a growth disadvantage. Likewise, decreasing ERK activity in cells expressing oncogenic RAS or RAF will allow cells to resume proliferation again. Indeed, hyperactivation of MAPK signaling is deleterious to BRAF^V600E^ melanoma cells (Leung et al., 2019). Moreover, melanomas with acquired resistance to RAF and MEK inhibitors become drug dependent for their continued proliferation due to elevation of their MAPK signaling (Thakur et al., 2013). On the contrary, in normal cells, reducing ERK activity rescued cells from senescence and facilitated cell transformation by oncogenic RAS without the need of additional oncogene expression (Deschênes-Simard et al., 2013). These results all pointed out a tumor-suppressive role of ERK signaling and support our bell-shaped ERK-proliferation model.

Our study suggests that the intensity of ERK signaling plays a pivotal role in determining the final proliferation outcome. Yet, how the strength of ERK signaling connects to the outputs of this pathway is largely unclear. Induction of BRAF^V600E^ for different amounts of time with and without treatment of cells with ERKi at different doses made it possible to generate cells with a wide range of ERK activities. RNA sequencing revealed the existence of four distinct classes of ERK-regulated genes which differed with respect to the relationship between ERK activity and the level of gene expression. The levels of Class I genes are linearly proportional (or inversely proportional) to ERK activity across the full range of ERK activity. Class II or III genes exhibit a convex relationship to ERK activity, and are most affected at either low or high ERK concentrations. Class IV genes exhibit a bell-shaped or U-shaped response, mimicking the response of proliferation to ERK activity. Previous studies that focused on gene expression at only high or low ERK levels would have been unable to resolve these 4 classes of ERK-dependencies. For example, genes that increase with ERK activity in all 4 classes would have been categorized in a single group when assayed at a high ERK level. The methods used here enable further refining of specific ERK-dependent signatures by identifying whether the genes are full-range linear ERK responders, high-range ERK responders or U-shape ERK responders. This analysis suggests that suppression of proliferation at low and high ERK levels likely proceeds via distinct transcriptional programs. Our ERK activity- and time-dependent classifications pave the way for dissecting the relevant pathways at different levels of ERK activity.

Our classification of genes also suggests that a single class of genes or a combination of multiple classes of genes could cooperate to reach the bell-shaped proliferation response. For instance, positive cell cycle regulators in a bell-shaped class or negative cell cycle regulators (e.g. p15INK4B) in a U-shaped class can, on their own or in combination, generate a bell-shaped proliferation response. Likewise, in the low-range ERK responder class, positive cell cycle regulators (e.g. G1 cyclins) were down-regulated and CDK inhibitors were upregulated at low ERK levels. This combination could explain the reduced proliferation at low ERK activity. It is thus reasonable to assume that cells take a network approach rather than a single-gene strategy to regulate proliferation across a range of ERK activity levels. It is also important to note that our gene classes are not exhaustive and that additional types of regulations (e.g., protein levels or post-translational modifications) must co-exist for a robust bell-shaped proliferation response. For instance, high ERK activity can prevent cell cycle progression by inducing degradation of key regulators (Deschênes-Simard et al., 2013), by modulating senescence-associated secretomes (Di Mitri and Alimonti, 2016) or by engaging homeostasis at tissue levels (Ruiz-Vega et al., 2020). We envision a broader set of genes and diverse regulatory mechanisms are needed for cells to engage a robust and coherent bell-shape proliferation response.

In summary, our studies provide a detailed and comprehensive map of the input-output relationship between ERK activity, proliferation response, and gene expression programs in non-transformed cells. In particular, the data provide an explanation for cell-to-cell heterogeneity in OIS induction in a nominally uniform population of proliferating cells. Our data help to explain the observed bell-shaped relationship between MAPK signaling and proliferation while also revealing substantial complexity in time and activity-dependent changes in gene expression. Such insights should improve our ability to study OIS *in vivo* and ultimately develop treatment regimens and therapeutics that exploit OIS to block the growth of human cancers.

## Supporting information

Key Resources Table

## ACKNOWLEDGMENTS

We thank S. Boswell and J. Lin for their help; members of the Lahav laboratory for comments, support, and ideas; the Nikon Imaging Center at HMS for support with live cell imaging and Harvard Research Computing for use of the O2 High Performance Compute Cluster. This research was supported by NIH grants GM116864 (GL) and P50-GM107618 (GL & PKS) and by NCI grant U54-CA225088 (PKS), Jane Coffin Child Memorial Fund for Medical Research fellowship to JYC, CONACyT/Fundacion Mexico en Harvard (404476), and Harvard Graduate Merit Fellowship to JR, Novartis Foundation fellowship to LG, and a HFSP grant LT000259/2019-L1 to FF.

## AUTHOR CONTRIBUTIONS

JYC conceived and designed the study; GL and PKS supervised the work. JYC generated reagents, established cell lines and performed experiments. JYC, CH, LG and FF performed RNA sequencing analysis. JYC, CH and JR analyzed the experimental data. CT and SLS assisted with live imaging analysis using EllipTrack. BP and HJG provided EKAREN5 reporter constructs. JYC, CH, AJ, PKS and GL wrote the manuscript with input from other authors.

## STAR METHODS

### EXPERIMENTAL MODEL AND SUBJECT DETAILS

#### Cell lines

Human retinal pigment epithelial (RPE) cells immortalized with human telomerase expression (RPE-hTERT, a kind gift from S.J. Elledge, Harvard Medical School) were grown in DMEM/F12 supplemented with 10% fetal bovine serum (FBS), 2mM L-Glutamine, Antibiotic-Antimycotic (100U/ml penicillin, 100μg/ml streptomycin and 250ng/ml Amphotericin B), and 50μg/ml hygromycin B.

#### Cell line construction

To establish RPE/tet-BRAF^V600E^-HA cell line, C-terminal HA-tagged BRAF^V600E^ construct was made by cloning the full-length BRAF^V600E^ expression cassette (Addgene plasmid # 15269) into a lentiviral backbone with tet-inducible promoter (Addgene plasmid # 41394, pLIX402). RPE cells were then infected with lentivirus carrying pLIX402-BRAF^V600E^-HA and selected with puromycin (2 g/ml) to obtain mixed cell clones. Single cell clones were expanded through limited dilution and subsequently screened for HA expression in the presence of doxycycline. To establish dE2F PIP reporter lines, expression cassette harboring *Drosophila* E2F1 PIP fragment (comprised of a.a. 1-187) fused to the C-terminus of Venus or mCherry fluorescent protein was cloned into the CSII-EF1 lentiviral vector. RPE cells transduced with lentiviruses carrying H2B-mTurquoise, mCherry-dE2F PIP and Venus-Geminin (1-110) or lentiviruses carrying H2B-mTurquoise and Venus-dE2F PIP were sorted on a BD FACSAria II high speed cell sorter to obtain pure populations expressing the desired fluorescent proteins. To establish RPE/EKAREV-NLS reporter line, RPE cells were co-transfected with pPB-CAG-EKAREV-NLS (Komatsu et al., 2011) and pCMV-hyPBase transposase vector (A. Bradley, Sanger Institute) and FACS sorted to obtain pure populations. To establish RPE/tet-BRAF^V600E^-HA + EKAREN5 + mCherry-dE2F PIP dual reporter line, verified RPE/tet-BRAF^V600E^-HA single cell clone was transduced with EKAREN5 (ERK FRET reporter) (Ponsioen et al., 2021) and mCherry-dE2F PIP lentiviruses and single cell clones harboring both reporters were obtained through single-cell sorting and subsequent expansion.

#### Time-lapse microscopy

Cells were plated in poly-D-lysine-coated glass-bottom plates (MatTek Corporation) and switched to phenol-red free culture medium supplemented with 10% FBS prior to live imaging. Cells were imaged using a Nikon Eclipse TE2000 microscope equipped with a chamber for controlled temperature (37%) and CO2 (5%) environment. All live-cell imaging was performed with a 10x Plan Apo objective (Nikon) and a Hamamatsu Orca ER camera using CFP, YFP, mCherry filter sets (Chroma). For EKAREV and EKAREN5 reporter imaging, the FRET signal was collected using customized ECFP/EYFP FRET filter sets with ET436/20x, ET535/30m, and T455lp mounting into the Nikon TE2000/Ti cube.

#### siRNA knockdown

Synthetic siRNAs used for this study were from Dharmacon siGenome SMART pool and were used at 13.3nM with Lipofectamine 2000 reagents (Invitrogen) according to manufacturer’s protocol. The following siRNAs were used: control siRNA (non-targeting #2), siGenome pooled set of four siRNAs for p16, p21 and p27. Specific antibodies were used to verify the target knockdown.

#### Western blot and senescence associated β-galactosidase assay

Cells were harvested in Laemmli Sample Buffer (Bio-Rad, #1610737) for 5 min at 95°C. Protein samples were separated by electrophoresis using 4-20% polyacrylamide gels (Bio-Rad, #456-9036) and transferred to Immun-Blot PVDF membranes (Bio-Rad #1620177). Blots were incubated with primary antibodies (p-RB: Cell Signaling #8516; p-ERK: Cell Signaling #4370; Actin: Sigma-Aldrich #5316; HA: Roche #11867423001) overnight at 4°C and then with HRP conjugated secondary antibodies for 1 hour at room temperature. HRP was detected using ECL substrates (Thermo Scientific #34076) and myECL Imager (Thermo Scientific).

#### Immunofluorescence

Immunostaining was performed in 96-well plates and all washes were done with the EL406™ Microplate Washer (BioTek). In brief, cells were fixed with 4% paraformaldehyde for 20 min, permeabilized with 0.2% Triton X-100 for 15 min and blocked with Odyssey® blocking buffer (LI-COR) for 1h before applying different antibodies. Primary antibodies were incubated overnight at 4°C. Appropriate Alexa Fluor® conjugated secondary antibodies were then used. For EdU staining, cells were pulsed with 10 M EdU for 30 min (or as indicated) prior to fixation and processed according to manufacturer’s instructions (Invitrogen #C10340). Cells were imaged with a 10x objective using an Operetta High Content Imaging System (Perkin Elmer, CT) or ImageXpress Micro Confocal High-Content Imaging System (Molecular Devices, CA). 9 sites were imaged in each well for 96-well plates.

#### Image analysis

Images for the immunostaining experiments were analyzed using MATLAB image analysis programs (Salmeen et al., 2010). Briefly, nuclear centroids were identified in images of Hoechst staining after applying a low-pass Gaussian filter and local background subtraction. A nucleus mask was generated for each cell by expansion from the centroid to reach 30% of maximum intensity. The nuclear pERK, BRAF^V600E^-HA, EdU and pRB mean intensity were measured after local background subtraction. The threshold level used to determine pRB and EdU positive cells was set using a k-means clustering algorithm on a day-to-day experiment basis.

#### Single-cell tracking and quantification following live-imaging

For population analysis of individual time frames, images were quantified and analyzed using MATLAB scripts (Cappell et al., 2016). To track single-cells following long-term live imaging, cells were tracked semi-automatically using a combined method of EllipTrack (Tian et al., 2020) and p53Cinema Single Cell Tracking Software (Reyes et al., 2018). In brief, EllipTrack segments cells by fitting nuclear contours with ellipses and tracks cells using a machine learning algorithm. The cell tracks were then manually curated using p53Cinema Single Cell Tracking Software that allows for real-time user correction of tracking and annotation of division events. Finally, verified tracks were kept for downstream analysis and signals from each color channel were extracted in the cell nuclei. G1-S and S-G2 transition events were computationally identified based on the Venus (or mCherry)-dE2F PIP intensity changes (1st derivative of the intensity) between frames. Briefly, Venus (or mCherry)-dE2F PIP levels would sharply decline or rise up when G1-S or S-G2 transition occurs, respectively.

#### ERK FRET reporter quantification

To quantify ERK activity, CFP, YFP and FRET images were acquired in RPE/ tet-BRAF^V600E^-HA + EKAREN5 + mCherry-dE2F PIP dual reporter cells. FRET images were taken by CFP excitation and YFP emission. Images were then subjected to flat field correction (to eliminate uneven illumination) and local background subtraction. The FRET signal was calculated on a pixel-by-pixel basis as follows. First, a FRET image was corrected for bleed-through from CFP and YFP channels.

[FRET]_corr_ = ([FRET]_Raw_-α [CFP] – β[YFP]
α: bleed-through of CFP into FRET channel upon CFP excitation
β: bleed-through of YFP into FRET channel upon CFP excitation of YFP

The two microscope-specific bleed-through parameters, α (0.53) and β (0.23), were determined using cells transfected with CFP or YFP alone. Then, the corrected FRET image ([FRET]corr) was normalized by the CFP image to obtain the FRET signal ([FRET]_corr_/[CFP]). ERK activity was calculated from the median value from the nuclear compartment of the FRET signal of each cell.

#### Statistical analysis

Error bars represent the standard deviation, standard error of the mean, or 95% bootstrap confidence interval as indicated in the legends. Statistical comparisons (p values) were obtained from two-sided t tests or otherwise as noted. The Pearson’s correlation coefficients (R) were calculated as indicated.

#### Total RNA sample preparation and quality control

RPE/tet-BRAF^V600E^-HA cells were plated in 6-well plates (75,000 per well) and allowed to grow for 72 hr till 50% confluency. Cells were then treated with the indicated concentrations of ERK inhibitor (SCH772984) alone for 24h or in combination with DOX (250ng/ml) for variable length of time as indicated by the experimental design. Each condition was performed twice on two different days for a total of two biological replicates. Cells were lysed with 600□l Trizol per well and total RNA was prepared using Direct-zol-96 RNA Kits according to the manufacturer’s protocol. Sample concentrations were determined by Nanodrop and RNA quality was assessed on a subset of samples by Bioanalyzer (Agilent); all samples scored RINs of > 9.0.

#### RNA sequencing library preparation

RNA sequencing library preparation was performed with the High Throughput TruSeq Stranded mRNA Library Prep Kit (Illumina) following the manufacturer’s protocol at half reaction volume. Input for each sample consisted of 300-500ng of RNA and 5μl of 1:500 diluted ERCC spike-in mix 1 (Ambion). Libraries were amplified for 12 cycles during the final amplification step. Libraries were quantified using the Qubit dsDNA HS assay (Thermo Fisher Scientific). Library size and quality were spot checked for a subset of samples by Bioanalyzer (Agilent). The average size of cDNA fragments in the libraries was 370 base pairs. Libraries were pooled at equimolar concentrations then the pool was quantitated using the KAPA library quantification kit (KAPA Biosystems). Libraries were sequenced single end 114 base pairs using NovaSeq_SP full flow cell (Illumina) at the Bauer Core Facility (Harvard University).

#### RNA-seq data processing

Reads were processed to counts using the bcbio-Nextgen toolkit (https://github.com/chapmanb/bcbio-nextgen) as follows: (1) Reads were trimmed and clipped for quality control in cutadapt v2.3; (2) Read quality was checked for each sample using FastQC v0.11.8; (3) High-quality reads were then aligned to the human assembly and gene annotation GRCh38.97 using Hisat2 v2.1.0 (Kim et al., 2019); (4) Genelevel transcript-counts were calculated using HTseq-count v0.9.1. Only data from genes annotated as protein-coding according to annotation from GRCh38.97 were kept for further analysis. Gene expression data (RNA seq) were deposited in the GEO (Gene Expression Omnibus, https://www.ncbi.nlm.nih.gov/geo/, accession number: GEO: GSE180210).

#### Differential expression analysis

Differential expression of genes was analyzed using DESeq2 (Love et al., 2014) by fitting a linear mixed effect that expressed the number of reads for each gene *K* using a negative binomial distribution of the form *K* = *NB*(*μ,σ*^2^) with mean *μ* and dispersion *σ*. The mean for each gene was modeled by a linear equation taking into account the sample treatments *T_x_* and batch with the untreated sample at time zero as intercept.

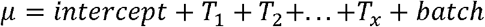

Log-fold changes were adjusted using procedures implemented in apeglm (Zhu et al., 2019), which estimates posterior distributions of the coefficients in the linear models that were fitted by DESeq2.

#### Selecting differentially expressed genes

In this setting, inferring differential expression poses a substantial challenge, since both dose-response and temporal dynamics need to be accounted for. To account for dependency on both time and dose, we used multiple regression with quadratic terms to describe the time-dose response surface (Log-fold changes) for every gene. As the same time points and doses were measured for all genes, we only computed a single QR-decomposition of the Vandermonde regressor matrix (Macon and Spitzbart, 1958) and then computed regression using matrix-matrix multiplication. To identify differential expression, we compared the goodness of fit between quadratic regression and constant approximation via the mean. P-values were computed using standard likelihood-ratio test and multiple-testing corrected using Bonferroni-Holm (Holm, 1979).

#### Clustering ERK-dependent differences in gene expression

Principal component analysis was performed on the matrix of moderated log2-fold changes of all samples compared to the baseline condition. The first principal component was strongly correlated with ERK activity across all samples. In order to find genes that have similar relationships between gene expression and ERK dose, we applied k-medoids clustering (Schubert and Rousseeuw, 2019) — a variation of the k-means clustering method that is robust to outliers — on the log2-fold changes at 24h. We found that choosing *k* = 8 resulted in easily interpretable clusters with different dynamics of ERK signaling responses. The clusters we identified can be grouped into four different response types, each with two variants representing responses with opposite signs.

#### Time series analysis

In order to group genes into clusters depending on their time of induction or repression, we first normalized the time series log2-fold changes of each gene. First, we computed the range *R* of log2-fold changes *f* for each gene *g* and each ERKi concentration *c* across all time points t by subtracting the minimum log2-fold change from the maximum.

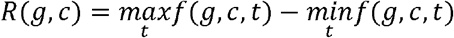

Next, we aggregated the time series data across all ERKi doses by computing 33% and 67% quantiles of log2-fold changes at each time point, weighted by the time series range *R*(*g,c*), using the algorithm *Q* described in (Harrel and Davis, 1982).

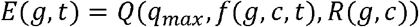

with 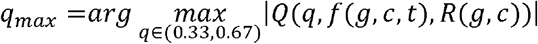

At each timepoint, the quantile with the highest absolute value was selected as the aggregate log2-fold change *E*, analogous to the procedure used for aggregating expression data across cell lines in the cited reference (Subramanian et al., 2017). Finally, genes were grouped into clusters based on when in the time series the absolute aggregated log2-fold change exceeded half of the maximum value across the entire time series. The significance of differences in the induction time between ERK-response clusters were tested using ANOVA followed by Tukey’s honest significant difference test. The distribution of the mean induction time for each ERK-response cluster was estimated using bootstrapping. The induction times of genes from each cluster were resampled 1000 times with replacement and the mean of each sample was computed.

#### Gene set enrichment analysis

Two variants of gene set enrichment analysis were performed. First, the enrichment of GO-terms in the ERK-response cluster gene sets was assessed using the R Bioconductor package topGO (https://bioconductor.org/packages/release/bioc/html/topGO.html). We considered all GO-terms in the Biological Process (“BP”) and Molecular Function (“MF”) categories. Enrichment was computed using the “weight01” algorithm and Fisher’s exact test. Second, gene set enrichment analysis was performed using gene sets from MSigDB (Liberzon et al., 2011). Specifically, gene sets from the Hallmark (H), curated pathways (C2:CP), and ontology (C5) categories, excluding Human Phenotype Ontology (HPO), were considered. We tested for significant enrichment using Fisher’s exact test on the overlap between the 1958 differentially expressed genes and the gene set of interest. Additionally, we computed the enrichment of a collection of manually curated gene sets related to ERK signaling, containing all gene sets containing “ERK”, “MAPK”, “senescence”, or “melanoma” from MSigDB, as well as the set of differentially expressed genes from a BRAF^V600E^ overexpression experiment (Capell et al., 2016) from GEO (GSE46801).

#### Replicate similarity

In order to assess the quality of replicates, we computed the Pearson correlation coefficients between the normalized counts of our two replicates, considering the 1000 most differentially expressed genes across all conditions. The genes were ranked by the results of a likelihood-ratio test using DESeq2, comparing the full model described above against a reduced model of the form *μ* = *intercept + batch*. The correlation matrix was plotted using the R package ComplexHeatmap (Gu et al., 2016).

