## Supplementary material for "Multi range ERK responses shape the proliferative trajectory of single cells following oncogene induced senescence": Key Resources Table

| REAGENT or RESOURCE | SOURCE | IDENTIFIER |
| --- | --- | --- |
| Antibodies | | |
| Phospho-p44/42 MAPK (Erk1/2) (Thr202/Tyr204) (D13.14.4E) antibody | Cell Signaling Technology | Cat# 4370, RRID: AB_2315112 |
| Phospho-Rb (Ser807/811) (D20B12) antibody | Cell Signaling Technology | Cat# 8516, RRID: AB_11178658 |
| Anti-β-Actin monoclonal antibody | Sigma-Aldrich | Cat# 5316, RRID: AB_476743 |
| Anti-Raf-B Antibody (F-7) | Santa Cruz Biotechnology | Cat# sc-5284, RRID: AB_626760 |
| Anti-HA High Affinity; Rat monoclonal antibody (clone 3F10) | Roche | Cat# 11867423001, RRID: AB_390918 |
| Goat anti-Rabbit IgG (H+L) Highly Cross-Adsorbed Secondary Antibody, Alexa Fluor 488 | Thermo Fisher Scientific | Cat# A-11034, RRID: AB_2576217 |
| Goat anti-Rat IgG (H+L) Cross-Adsorbed Secondary Antibody, Alexa Fluor 568 | Thermo Fisher Scientific | Cat# A-11077, RRID: AB_2534121 |
| Goat anti-Mouse IgG (H+L) Cross-Adsorbed Secondary Antibody, Alexa Fluor 647 | Thermo Fisher Scientific | Cat# A-21235, RRID: AB_2535804 |
| Bacterial and virus strains | | |
| CSII-EF1-H2B-mTurquoise (lentiviral) | (Spencer et al., 2013) | N/A |
| CSII-EF1-mVenus-hGeminin(1–110) (lentiviral) | (Sakaue-Sawano et al., 2008) | N/A |
| CSII-EF1-mCherry-dE2F PIP (lentiviral) | This work | N/A |
| CSII-EF1-mVenus-dE2F PIP (lentiviral) | This work | N/A |
| LV-EKAREN5-NLS (lentiviral) | Addgene | Plasmid#167818 |
| LIX402-BRAF^V600E^-HA-Puro (lentiviral) | This work | N/A |
| Biological samples | | |
| Chemicals, peptides, and recombinant proteins | | |
| Hoeschst 33342, Trihydrochloride, Trihydrate | Thermo Fisher Scientific | Cat# H3570 |
| SCH772984, ERK inhibitor | MedChem Express | Cat# HY-50846 |
| Doxycycline hyclate | Sigma-Aldrich | Cat# D9891-5G |
| SMARTpool: siGENOME Non-targeting siRNA control pools | Horizon Discovery | Cat# D-001206-14-05 |
| SMARTpool: siGENOME Human CDKN1A siRNA | Horizon Discovery | Cat# M-003471-00-0005 |
| SMARTpool: siGENOME Human CDKN2A siRNA | Horizon Discovery | Cat# M-011007-03-0005 |
| SMARTpool: siGENOME Human CDKN1B siRNA | Horizon Discovery | Cat# M-003472-00-0005 |
| Critical commercial assays | | |
| Lipofectamine 2000 Transfection Reagent | Thermo Fisher Scientific | Cat# 11668027 |
| Senescence Associated 𝛽-Galactosidase staining kit | Cell Signaling Technology | Cat# 9860 |
| Click-iT™ Cell Reaction Buffer Kit | Thermo Fisher Scientific | Cat# C10269 |
| Click-iT™ EdU (5-ethynyl-2’-deoxyuridine) | Thermo Fisher Scientific | Cat# A10044 |
| Click-iT™ Alexa Fluor® 647 Azide, Triethylammonium Salt | Thermo Fisher Scientific | Cat# A10277 |
| Deposited data | | |
| Raw and analyzed RNA-seq data | GEO (Gene Expression Omnibus) | GEO: GSE180210 |
| Processed data and results used to generate main and supplemental figures in the manuscript | Synapse database | <https://www.synapse.org/#!Synapse:syn21411369> |
| Experimental models: Cell lines | | |
| Human: RPE hTERT | S.J. Elledge Lab (Harvard Medical School) | N/A |
| Human: RPE + tet-BRAF^V600E^-HA | This work | N/A |
| Human: RPE + H2B-mTurquoise + Venus-dE2F PIP | This work | N/A |
| Human: RPE + tet-BRAF^V600E^-HA + EKAREN5 + mCherry-dE2F PIP | This work | N/A |
| Human: RPE + EKAREV-NLS | This work | N/A |
| Experimental models: Organisms/strains | | |
| Oligonucleotides | | |
| Recombinant DNA | | |
| pPB-CAG-EKAREV-NLS (piggyBac) | (Komatsu et al., 2011) | N/A |
| pCMV-hyPBase | (Komatsu et al., 2011) | N/A |
| Software and algorithms | | |
| Scripts for analysis of RNA sequencing data | This work | <https://github.com/clemenshug/erk_senescence> |
| Software for automatic segmentation and quantification of immunofluorescence images | (Salmeen et al., 2010) | N/A |
| Software for automatic segmentation, and quantification of fluorescent reporter cells following live imaging | (Cappell et al., 2016) | <https://github.com/scappell/Cell_tracking> |
| p53 Cinema Single Cell Tracking | (Reyes et al., 2018) | <https://github.com/balvahal/p53CinemaManual> |
| EllipTrack | (Tian et al., 2020) | <https://github.com/tianchengzhe/elliptrack> |
| Other | | |
