## Supplemental Figures for "Multi range ERK responses shape the proliferative trajectory of single cells following oncogene induced senescence"

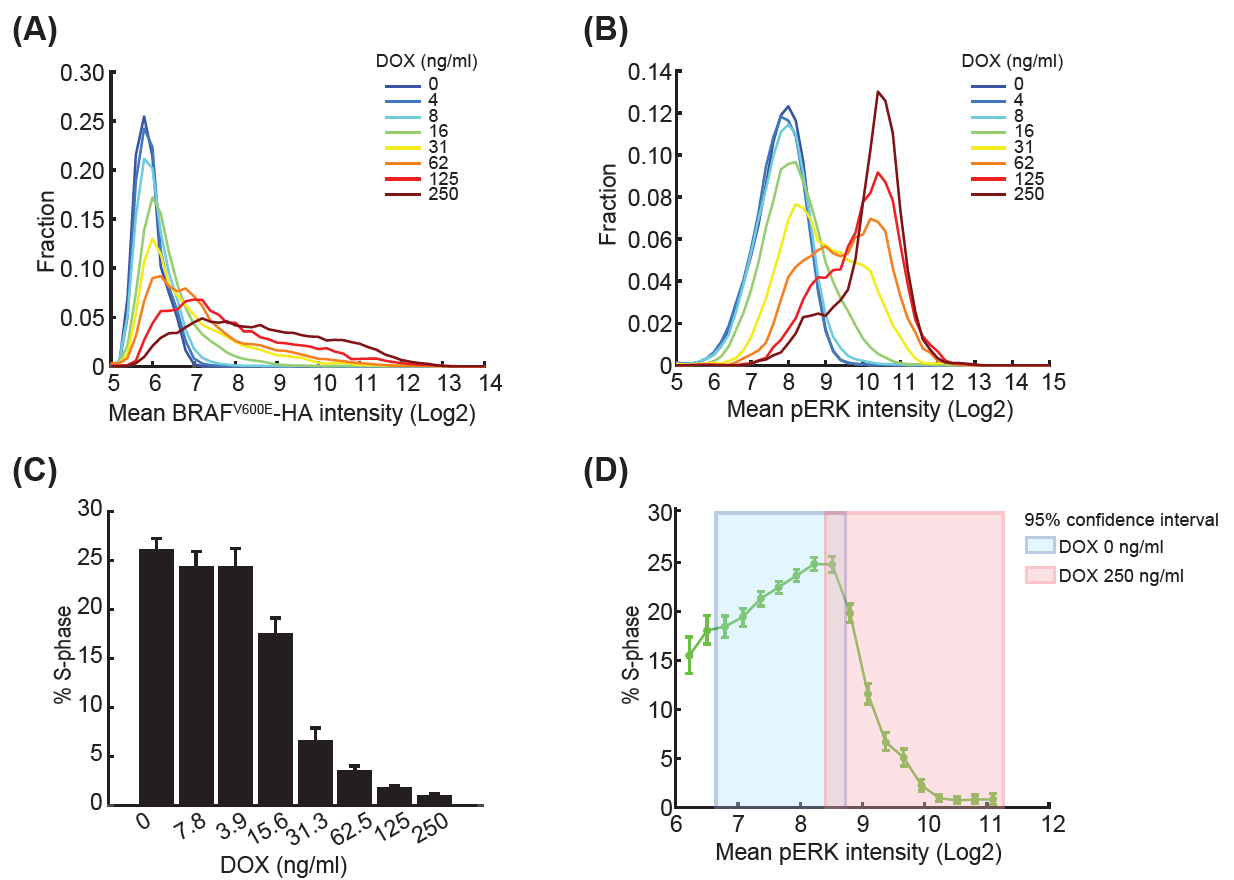

### Figure S1. Tunable expression of BRAF^V600E^ combined with heterogeneity in ERK activation and proliferation achieves a wide range of downstream responses.

**(A & B)** Frequency plots showing BRAF^V600E^ (A) and pERK (B) distribution following DOX treatment. RPE/tet-BRAF^V600E^-HA cells were treated with serial doses of DOX (0-250ng/ml, 2-fold dilution) for 72 hr and immunostained for HA and pERK.

**(C)** As in (A & B), RPE/tet-BRAFV600E-HA cells were pulsed with EdU for 30 min after 72 h of DOX treatment and the percent in S phase (%S) were quantified and plotted as a function of the DOX dose.

**(D)** Ranges of proliferation (percent of cells in S phase) observed at lower pERK levels in the absence of DOX or at higher pERK levels induced by DOX (250ng/ml) 72h post treatment. Shaded area shows a 95% confidence interval of pERK intensity treated with 0 or 250ng/ml DOX.

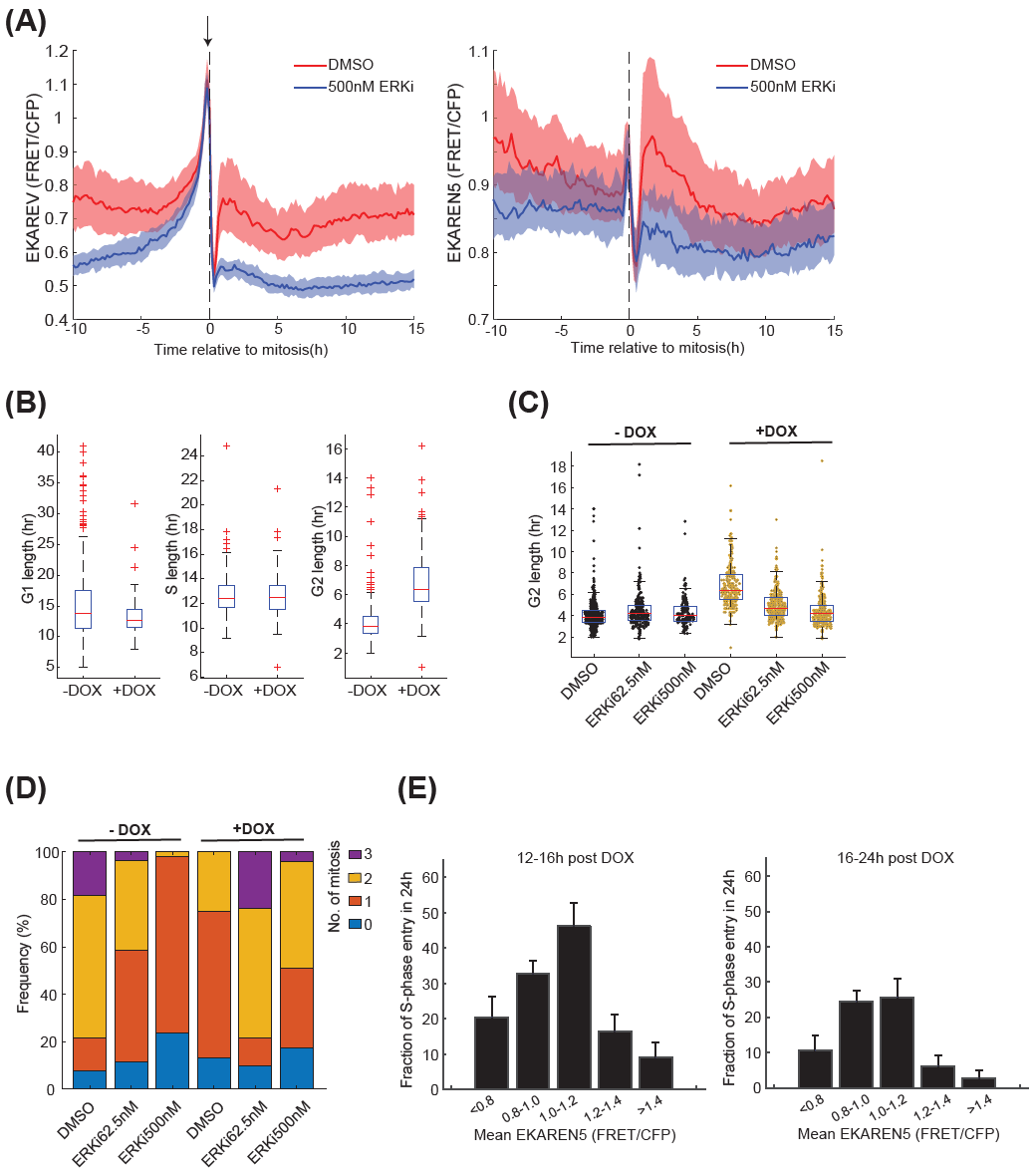

### Figure S2. ERK activation lengthens G2 duration and decreases cell divisions.

**(A)** Dynamics of ERK activity measured by the traditional EKAREV (left) or the improved EKAREN5 probe (right) in the vicinity of mitosis. RPE cells stably expressing the EKAREV reporter or the EKAREN5 reporter were treated with DMSO or 500nM ERK inhibitor followed by 48h of live imaging (EKAREV DMSO, n=294; EKAREV ERKi, n=100; EKAREN5 DMSO, n=139; EKAREN5 ERKi, n=48). Cells were *in silico* synchronized at mitosis. Bold lines and shaded areas correspond to median and interquartile range, respectively. Non-specific activity of EKAREV prior to mitosis is indicated with an arrow.

**(B)** Box plots comparing G1 (left), S (middle) and G2 (right) duration in untreated or DOX-treated RPE/tet-BRAF^V600E^-HA + EKAREN5 + mCherry-dE2F PIP Dual Reporter cells. Duration of each cell cycle phase in individual cells was computationally derived based on the mCherry-dE2F PIP reporter. n=260 (-DOX, G1), 61 (+DOX, G1), 334 (-DOX, S), 164 (+DOX, S), 399 (-DOX, G2), and 236 (+DOX, G2).

**(C)** Box plots comparing G2 duration in RPE Dual Reporter cells treated with DMSO, 62.5nM ERKi or 500nM ERKi in the absence or presence of DOX. Each dot represents a single cell (n>130 cells per condition).

**(D)** Frequency of divisions in RPE Dual Reporter cells treated with DMSO, 62.5nM ERKi or 500nM ERKi in the absence or presence of DOX. Individual cells were tracked for 72h; after division, one daughter cell was randomly selected for further tracking (also see **Fig 3B**). n>200 cells for each condition.

**(E)** Fraction of S-phase entry in response to increasing ERK activity. Similar to **Fig 3D**, data from all treatments were pooled together and the mean ERK activity between 12-16h and 16-24h post treatment was calculated. The probability of entering into S-phase was quantified within 24h after the time-frame in which ERK activity was monitored (mean ± 95% confidence interval, n>150 for each binning ERK activity).

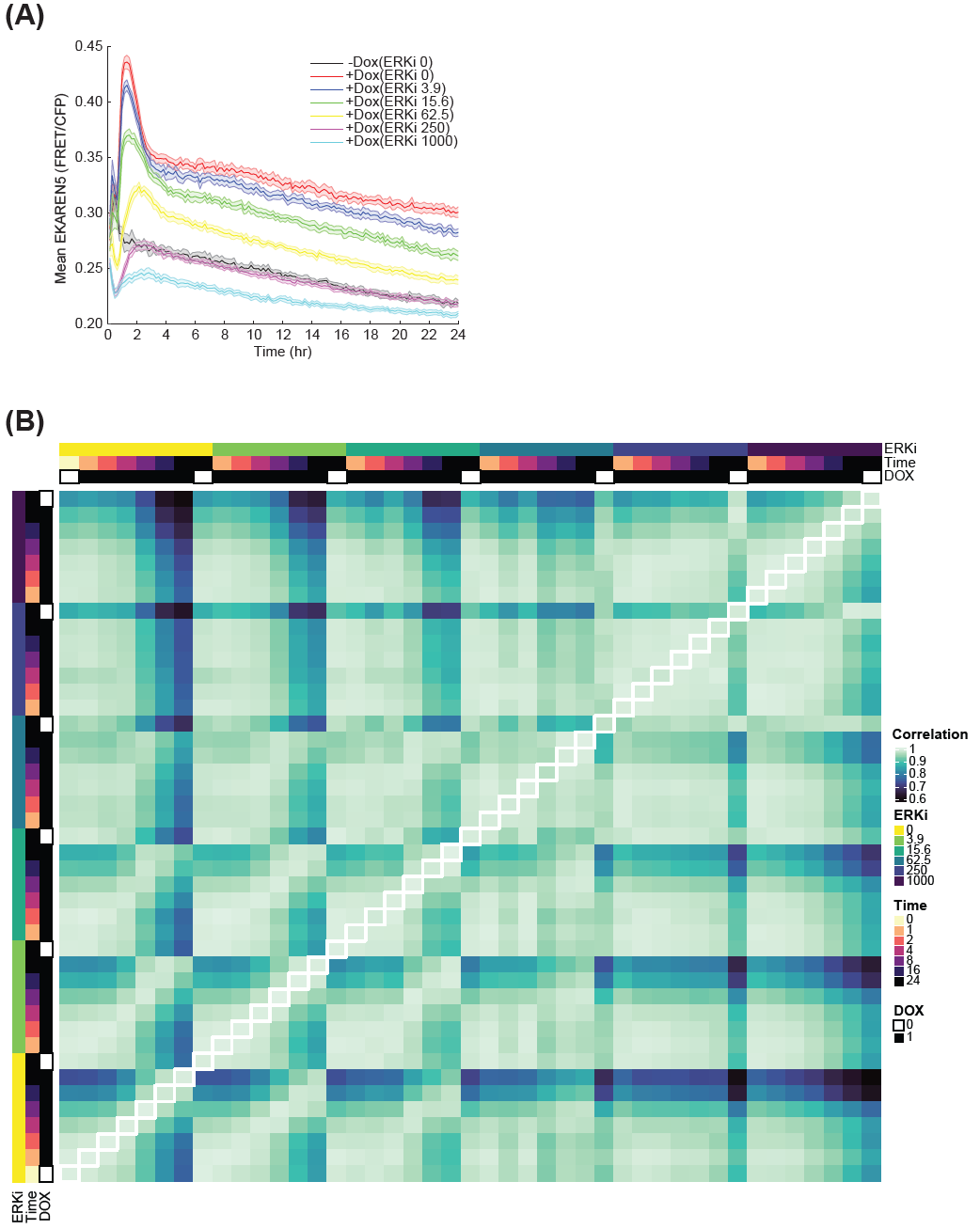

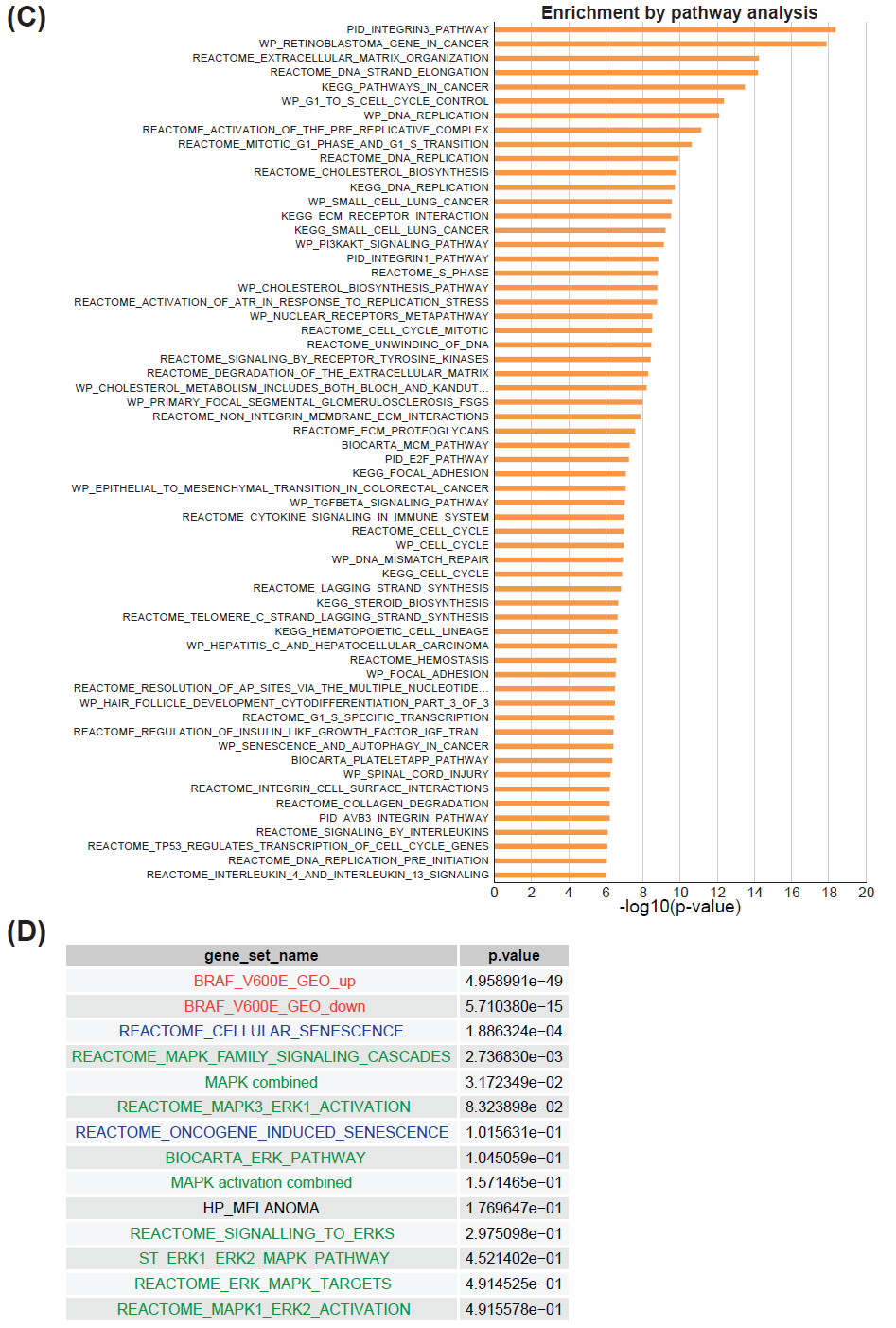

### Figure S3. ERK activity and pairwise correlations between RNA-seq replicates as a function of time and ERK inhibitor dose, and gene set enrichment analysis of ERK-responsive genes.

**(A)** Dynamics of ERK activity following DOX and ERK inhibitor treatments. RPE Dual Reporter cells were treated with DMSO (ERKi 0) in the absence or presence of DOX or with different concentrations of ERK inhibitor in the presence of DOX followed by 24 hr of live imaging (median ± SD of fourteen imaging positions).

**(B)** Heatmap of pairwise Pearson correlation coefficients between all RNA-seq replicates, with one replicate on the x-axis and the other on the y-axis. Replicates with the same treatments are marked with white stroke. In the margins of the heatmap, the treatment of samples with ERK inhibitor, doxycycline, and time of collection are shown. Correlations are computed using normalized gene counts from DESeq2, considering only the 1000 most differentially expressed genes.

**(C)** Gene set enrichment analysis (GSEA) of the 1958 genes that were significantly differentially expressed across all conditions (see **Fig 4E**). Shown are the most enriched MSigDB gene sets in the Hallmark (H), curated pathways (C2: CP), and ontology (C5) categories, excluding Human Phenotype Ontology (HPO).

**(D)** GSEA of selected BRAF^V600E^ and ERK-related gene sets amongst 1958 significantly expressed genes. ERK-related gene sets include all gene sets in MSigDB that mentioned “ERK”, “MAPK”, “senescence”, or “melanoma”. BRAF^V600E^ related gene sets include the set of differentially expressed genes from a BRAF^V600E^ overexpression experiment (Capell et al., 2016) from GEO (GSE46801).

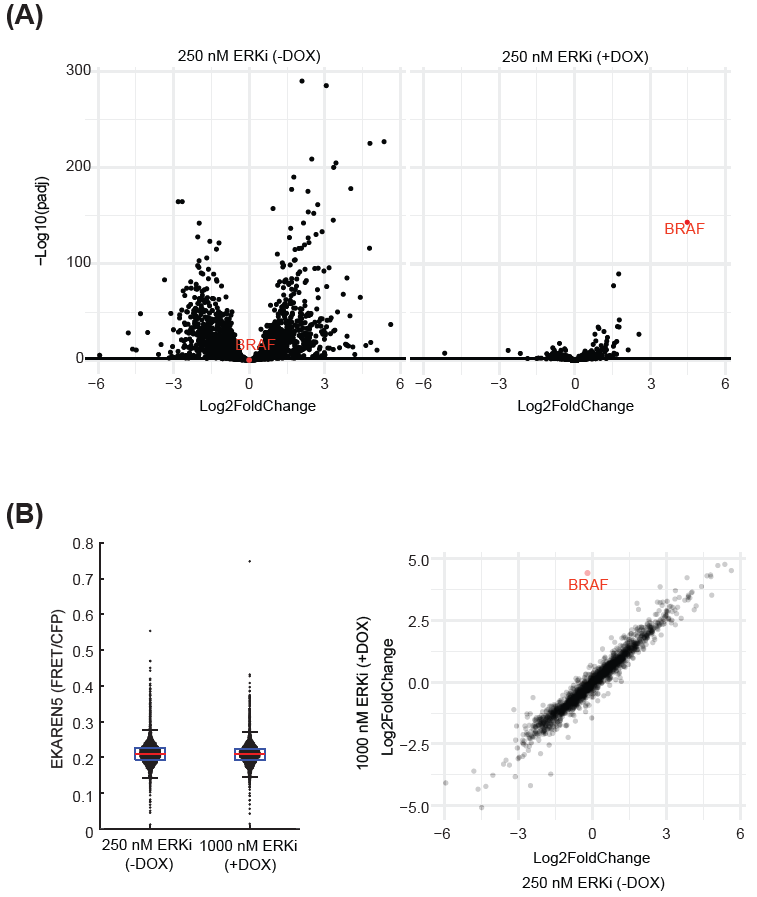

### Figure S4 Rescue of ERKi effects on gene expression by BRAF^V600E^ overexpression.

(A) Volcano plots of the differential gene expression between two treatment conditions and the untreated control condition. On the left, inhibition with 250 nM ERKi results in large changes in gene expression (438 genes; absolute log2FoldChange > 1; p < 0.05). On the right, in addition to 250 nM ERKi, doxycycline was used to induce BRAF^V600E^ overexpression (highlighted in red). At this concentration of ERKi, its effects were largely rescued by BRAF^V600E^ overexpression, with only 42 differentially expressed genes remaining.

(B) (Left) Box plots showing similar ERK activities in BRAF^V600E^ Dual Reporter cells treated with 250 nM ERKi in the absence of DOX, or 1000 nM ERKi in the presence of DOX. The ERK activity of each cell was measured at 24 hr post treatment. Each dot represents a single cell (n>5000 cells per condition). (Right) Correlation between the differential gene expression compared to untreated control of cells treated with 250 nM ERKi (x-axis) and 1000 nM ERKi plus doxycycline to induce overexpression of BRAF^V600E^ (y-axis). The two conditions display extremely similar differential gene expression (R-squared 0.97; p < 2.2e-16), indicating that the effect of BRAF^V600E^ overexpression can be rescued by increasing the concentration of the ERK inhibitor.

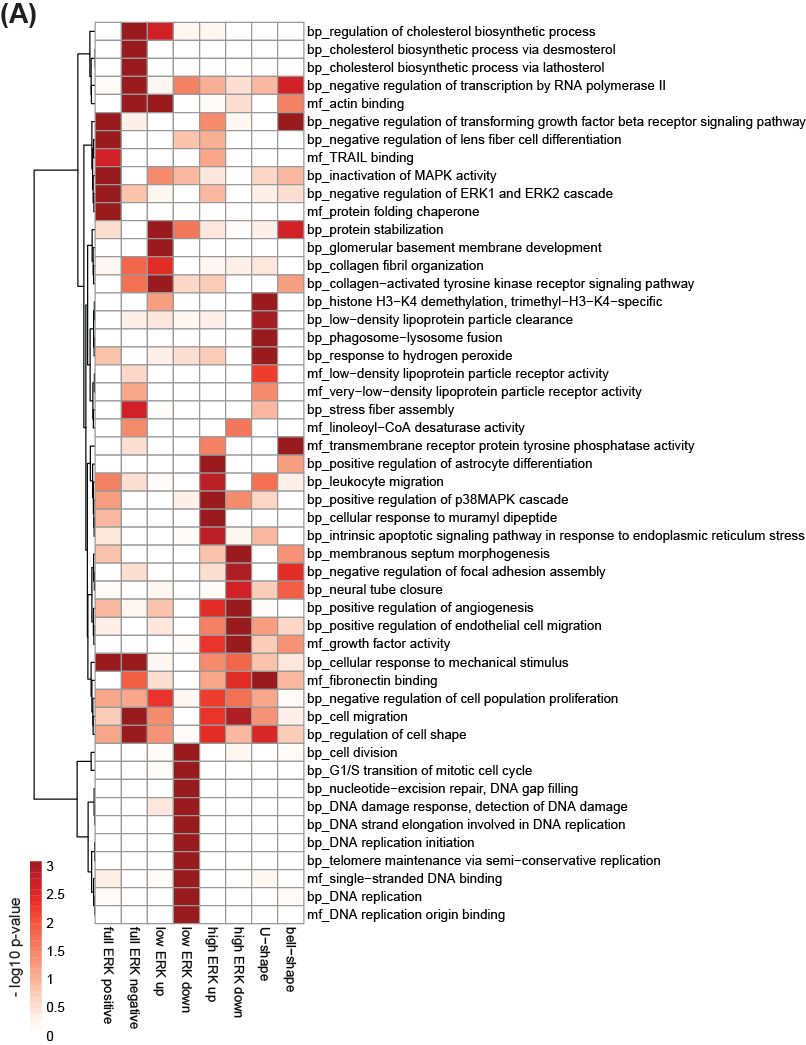

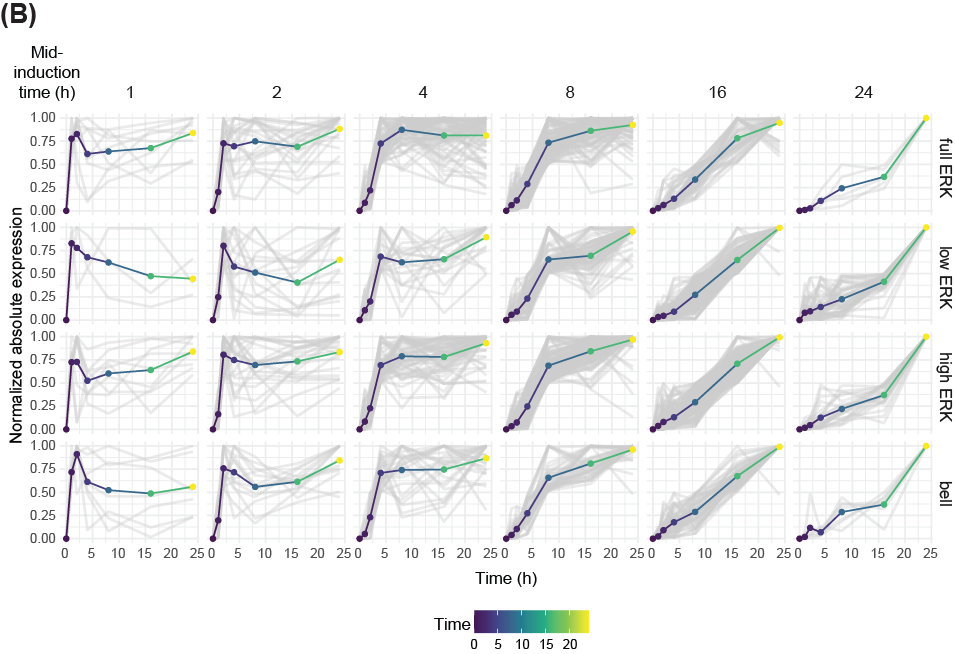

### Figure S5. Association of ERK responsive clusters with functional GO enrichment groups.

**(A)** GO term enrichment analysis for genes in all eight ERK response clusters. The 10 most significant GO terms per cluster are shown, with a maximum of 50 clusters in total. Rows are ordered using hierarchical clustering. Terms are prefixed with their GO domain. “mf” corresponds to Molecular Function and “bp” to Biological Function.

**(B)** Trajectory of gene expression over time. Gray lines represent normalized expression of individual genes, and colored lines show the average trajectory for the group. Each row represents an ERK-response cluster. Each column represents the time point at which genes in that group first reached half of their maximum induction. Vertically, the panels are divided into the four ERK response clusters that we identified. Gene expression was normalized to the maximum value observed in the dataset, and plotted as an absolute value ranging from 0 to 1.
